# Differential response of macrophages and neutrophils to trypanosome infections in zebrafish: occurrence of foamy macrophages

**DOI:** 10.1101/2020.11.01.363721

**Authors:** Sem H. Jacobs, Éva Dóró, Ffion R. Hammond, Mai E. Nguyen-Chi, Georges Lutfalla, Geert F. Wiegertjes, Maria Forlenza

## Abstract

A tightly regulated innate immune response to trypanosome infections is critical to strike a balance between parasite control and inflammation-associated pathology. In the present study, we make use of the recently established *Trypanosoma carassii* infection model in larval zebrafish to study the early response of macrophages and neutrophils to trypanosome infections *in vivo*. We consistently identified high- and low-infected individuals and were able to simultaneously characterize their differential innate response. Not only did macrophage and neutrophil number and distribution differ between the two groups, but also macrophage morphology and activation state. Exclusive to high-infected zebrafish, was the appearance of macrophages rich in lipid droplets, confirmed to be foamy macrophages and characterized by a strong pro-inflammatory profile. Altogether, we provide an *in vivo* characterization of the differential response of macrophage and neutrophil to trypanosome infection and identify foamy macrophages as potentially associated with an exacerbated immune response and susceptibility to the infection. To our knowledge this is the first report of the occurrence of foamy macrophages during an extracellular trypanosome infection.

## Introduction

Trypanosomes of the *Trypanosoma* genus are protozoan haemoflagellates that can infect animals from all vertebrate classes, including warm-blooded mammals and birds as well as cold-blooded amphibians, reptiles and fish. This genus contains human and animal pathogens, including the intracellular *Trypanosoma cruzi* (causing Human American Trypanosomiasis or Chagas’ disease), the extracellular *T. brucei rhodesiense* and *T. brucei gambiense* (causing Human African Trypanosomiasis or Sleeping Sickness) and *T. congolense, T. vivax* and *T. b. brucei* (causing Animal African Trypanosomiasis or Nagana) (Radwanska et al., 2018; Simpson et al., 2006). Among these, salivarian trypanosomes such as *T. brucei* ssp. live extracellularly in the bloodstream or tissue fluids of their host. For example, *T. vivax* can multiply rapidly and is evenly distributed throughout the cardiovascular system, *T. congolense* tends to aggregate in small blood vessels, whereas *T. brucei* especially can extravasate and multiply in interstitial tissues (reviewed by Magez and Caljon, 2011). Pathologically, anaemia appears to be a factor common to infections with most if not all trypanosomes although with different underlying causative mechanisms. These include, among others, erythrophagocytosis by macrophages (Cnops et al., 2015; Guegan et al., 2013), hemodilution (Naessens, 2006), erythrolysis through intermembrane transfer of variant surface glycoprotein (VSG) from trypanosomes to erythrocytes (Rifkin and Landsberger, 1990), oxidative stress from free radicals (Mishra et al., 2017) and mechanical damage through direct interaction of trypanosomes with erythrocytes surface (Boada-Sucre et al., 2016).

Immunologically, infections with trypanosomes are often associated with dysfunction and pathology related to exacerbated innate and adaptive immune responses (reviewed by Radwanska et al., 2018; Stijlemans et al., 2016). Initially it was believed that antibody-dependent complement-mediated lysis was the major protective mechanism involved in early parasite control (Krettli et al., 1979; Musoke and Barbet, 1977). However, later studies revealed that at low antibody levels, trypanosomes can efficiently remove surface-bound antibodies through an endocytosis-mediated mechanisms (Engstler et al., 2007), and that complement C5-deficient mice are able to control the first-peak parasitaemia similarly to wild type mice (La Greca et al., 2014). Instead, innate immune mediators such as IFNγ, TNFα and nitric oxide (NO) were shown to be indispensable for the control of first-peak parasitaemia, through direct and indirect mechanisms (reviewed by Radwanska et al., 2018). In the early phase of infection, the timely induction of IFNγ by NK, NKT and CD8^+^ cells (Cnops et al., 2015) followed by the production of TNFα and NO by IFNγ-primed macrophages (Baral et al., 2007; Iraqi et al., 2001; Lopez et al., 2008; Rudolf Lucas et al., 1994; Magez et al., 1993, 2007, 2006, 2001, 1999; O’Gorman et al., 2006; Sternberg and Mabbott, 1996; Wu et al., 2017) leads to effective control of first-peak parasitaemia. Glycosyl-inositol-phosphate soluble variant surface glycoproteins (GPI-VSG) released from the surface of trypanosomes were found to be the major inducers of TNFα in macrophages, and that such response could be primed by IFNγ (Coller et al., 2003; Magez et al., 2002). When macrophages would encounter GPI-VSG prior to IFNγ exposure however, their TNFα and NO response would dramatically be reduced (Coller et al., 2003) which, depending on the timing, could either lead to macrophage unresponsiveness or prevent exacerbated inflammatory responses during the first-peak of parasite clearance. Altogether, these data made clear that an early innate immune response is crucial to control the acute phase of trypanosome infection, but that its tight regulation is critical to ensure parasite control as opposed to pathology.

All the findings above took advantage of the availability of several mice models for trypanosome infection using trypanosusceptible or trypanotolerant as well as mutant ‘knock-out’ mice strains. Although mice cannot be considered natural hosts of trypanosomes and do not always recapitulate all features of natural infections, the availability of such models allowed to gain insights into the general biology of trypanosomes, their interaction with and evasion of the host immune system, as well as into various aspects related to vaccine failure, antigenic variation, and (uncontrolled) inflammation (Magez and Caljon, 2011). The use of knock-out strains for example, shed specific light on the role of various cytokines, particularly TNFα, IFNγ and IL-10, in the control of parasitaemia and in the induction of pathological conditions during infection (reviewed in Magez and Caljon, 2011). It would be ideal to be able to follow, *in vivo*, the early host responses to the infection and visualise the trypanosome response to the host’s attack. However, due to the lack of transparency of most mammalian hosts, this has not yet been feasible.

We recently reported the establishment of an experimental trypanosome infection of zebrafish (*Danio rerio*) with the fish-specific trypanosome *Trypanosoma carassii* (Dóró et al., 2019). In the latter study, by combining *T. carassii* infection of transparent zebrafish with high-resolution high-speed microscopy, we were able to describe in detail the swimming behaviour of trypanosomes *in vivo*, in the natural environment of blood and tissues of a live vertebrate host. This led to the discovery of novel attachment mechanisms as well as trypanosome swimming behaviours that otherwise would not have been observed *in vitro* (Dóró et al., 2019). Previous studies in common carp (*Cyprinus carpio*), goldfish (*Carrassius aurata*) and more recently zebrafish, demonstrated that infections with *T. carassii* present many of the pathological features observed during human or animal trypanosomiasis, including a pro-inflammatory response during first-peak parasitaemia (Kovacevic et al., 2015; Oladiran et al., 2011; Oladiran and Belosevic, 2009) polyclonal B and T cell activation (Joerink et al., 2007, 2004; Lischke et al., 2000; Ribeiro et al., 2010; Woo and Ardelli, 2014) and anaemia (Dóró et al., 2019; Islam and Woo, 1991; McAllister et al., 2019). These shared features among human and animal (including fish) trypanosomiases suggest a commonality in (innate) immune responses to trypanosomes across different vertebrates.

Zebrafish are fresh water cyprinid fish closely related to many of the natural hosts of *T. carassii* (Kent et al., 1993; Simpson et al., 2006) and are a powerful model species owing to, among others, their genetic tractability, large number of transgenic lines marking several immune cell types, knock-out mutant lines and most importantly, the transparency of developing embryos allowing high-resolution *in vivo* visualisation of cell behaviour (Benard et al., 2015; Bertrand et al., 2010; Ellett et al., 2011; Langenau et al., 2004; Lawson and Weinstein, 2002; Page et al., 2013; Petrie-Hanson et al., 2009; Renshaw et al., 2006; White et al., 2008). During the first 2-3 weeks of development, zebrafish are devoid of mature T and B lymphocytes and thus offer a window of opportunity to study innate immune responses (Torraca et al., 2014; Torraca and Mostowy, 2018), especially those driven by neutrophils and macrophages. The response of macrophages and neutrophilic granulocytes towards several viral, fungal and bacterial pathogens has been studied in detail using zebrafish (Cronan and Tobin, 2014; García-Valtanen et al., 2017; Nguyen-Chi et al., 2014a; Palha et al., 2013; Ramakrishnan, 2013; Renshaw and Trede, 2012; Rosowski et al., 2018; Torraca and Mostowy, 2018) but never before in the context of trypanosome infections.

Taking advantage of the recently established zebrafish-*T. carassii* infection model and of the availability of zebrafish transgenic lines marking macrophages and neutrophils as well as *il1b*- and *tnfa*-expressing cells, in the current study, we describe the early events of the innate immune response of zebrafish to *T. carassii* infections. Based on a novel clinical scoring system relying, amongst other criteria, on *in vivo* real-time monitoring of parasitaemia, we could consistently segregate larvae in high- and low-infected individuals without having to sacrifice the larvae. Between these individuals we always observed a marked differential response between macrophages and neutrophils, especially with respect to their proliferative capacity and redistribution in tissues or major blood vessels during infection. Significant differences were observed in the inflammatory response of macrophages in high- and low-infected individuals and in their susceptibility to the infection. In low-infected individuals, despite an early increase in macrophage number, a mild inflammatory response strongly associated with control of parasitaemia and survival to the infection was observed. Conversely, exclusively in high-infected individuals, we describe the occurrence of large, granular macrophages, reminiscent of foamy macrophages (Vallochi et al., 2018), characterized by a strong inflammatory profile and association to susceptibility to the infection. This is the first report of the occurrence of foamy macrophages during an extracellular trypanosome infection.

## Materials and methods

### Zebrafish lines and maintenance

Zebrafish were kept and handled according to the Zebrafish Book (zfin.org) and local animal welfare regulations of The Netherlands. Zebrafish embryo and larvae were raised in egg water (0.6 g/L sea salt, Sera Marin, Heinsberg, Germany) at 27°C with a 12:12 light-dark cycle. From 5 days post fertilisation (dpf) until 14 dpf larvae were fed Tetrahymena once a day. From 10 dpf larvae were also daily fed dry food ZM-100 (ZM systems, UK). The following zebrafish lines used in this study: transgenic *Tg(mpx:GFP)^i114^* (Renshaw et al., 2006), *Tg(kdrl:hras-mCherry)^s896^ referred as Tg(kdrl:caax-mCherry)* (Chi et al., 2008), *Tg(fli1:eGFP)^y1^* (Lawson and Weinstein, 2002), *Tg(mpeg1:eGFP)^gl22^* (Ellett et al., 2011)*, Tg(mpeg1.4:mCherry-F)^ump2Tg^, Tg(il1b:eGFP-F) ^ump3Tg^*, (Nguyen-Chi et al., 2014b), *Tg(tnfa:eGFP-F)^ump5Tg^* (Nguyen-Chi et al., 2015) or crosses thereof. The latter three transgenic zebrafish lines express a farnesylated (membrane-bound) mCherry (mCherry-F) or eGFP (eGFP-F) under the control of the *mpeg1, il1b* or *tnfa* promoter, respectively.

### *Trypanosoma carassii* culture and infection of zebrafish larvae

*Trypanosoma carassii* (strain TsCc-NEM) was cloned and characterized previously (Overath et al., 1998) and maintained in our laboratory by syringe passage through common carp (*Cyprinus carpio*) as described previously (Dóró et al., 2019). Blood was drawn from infected carp and kept at 4°C overnight in siliconized tubes. Trypanosomes enriched at the interface between the red blood cells and plasma were collected and centrifuged at 800 xg for 8 min at room temperature. Trypanosomes were resuspended at a density of 5 x 10^5^-1 x 10^6^ ml and cultured in 75 or 165 cm^2^ flasks at 27°C without CO_2_ in complete medium as described previously (Dóró et al., 2019). *T. carassii* were kept at a density below 5 x 10^6^/ml, and sub-cultured 1-3 times a week. In this way *T. carassii* could be kept in culture without losing infectivity for up to 2 months. The majority of carp white blood cells present in the enriched trypanosome fraction immediately after isolation, died within the first 3-5 days of culture and any remaining blood cell was removed prior to *T. carassii* injection into zebrafish. To this end, cells were centrifuged at 800 xg for 5 min in a 50 ml Falcon tube and the tube was subsequently tilted in a 20° angle in relation to the table surface, facilitating the separation of the motile trypanosomes along the conical part of the tube from the static pellet of white blood cells at the bottom of the tube.

For zebrafish infection, trypanosomes were cultured for 1 week and no longer than 3 weeks. Infection of zebrafish larvae was performed as described previously (Dóró et al., 2019). Briefly, prior to injection, 5 days post fertilization (dpf) zebrafish larvae were anaesthetized with 0.017% Ethyl 3-aminobenzoate methanesulfonate (MS-222, Tricaine, Sigma-Aldrich) in egg water. *T. carassii* were resuspended in 2% polyvinylpyrrolidone (PVP, Sigma-Aldrich) and injected (n=200) intravenously in the Duct of Cuvier.

### Clinical scoring system of the severity of infection

Careful monitoring of the swimming behaviour of zebrafish larvae after infection (5 dpf onwards) as well as *in vivo* observation of parasitaemia development in transparent larvae, led to the observation that from 4 days post infection (dpi) onwards larvae could generally be segregated into high- and low-infected individuals. To objectively assign zebrafish to either one of these two groups, we developed a clinical scoring system (**Fig 1**). The first criterion looked at the escape reflex upon contact with a pipette and was sufficient to identify high-infected individuals as those not reacting to the pipet (slow swimmers). To categorize the remaining individuals, an infection score based on counting parasite:blood cell ratios in 100 events passing through the intersegmental capillary (ISC) above the cloaca was developed. The infection scores on a scale from 1 to 10 were assigned as follows: 1=no parasites observed, 2=1-10% parasite, 3=11-20% parasite, 4=21-30% parasite, 5=31-40% parasite, 6=41-55% parasite, 7=56-70% parasite, 8=71-85% parasite, 9=86-99%, 10=no blood cells observed. Larvae with infection scores between 1-3 were categorized as low-infected while scores between 6-10 were categorized as high infected. Larvae with scores 4-5 were reassessed 1 day later, at 5 dpi, and then categorised as high- or low-infected. Next to that, the swimming behaviour of larvae was observed and compared to the control group. Heartbeat of the larvae was monitored and noted if it was slower than the control. The diameter of the cardinal caudal vein in the trunk area after the cloaca region was measured in ImageJ 1.49o to quantify the degree of vasodilation. Eventual blockage of tail tip vessel-loop was also noted. Extravasation and the location of extravasated parasites (e.g. fins, muscle, intraperitoneal cavity, and interstitial space lining the blood vessels) was recorded.

**Fig 1.**
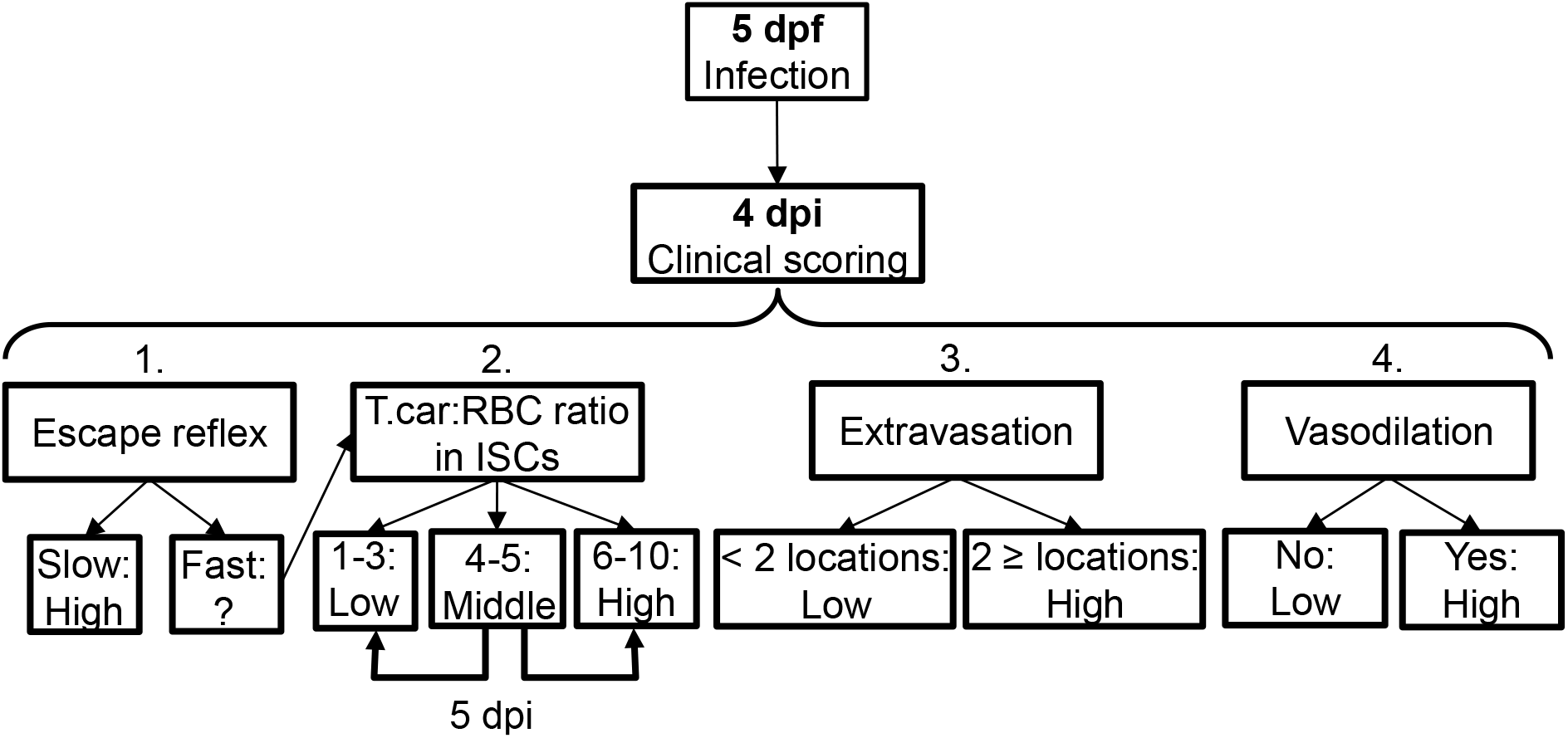
Schematic overview of the clinical scoring system used to determine individual infection levels of *T. carassii*-infected zebrafish larvae. Zebrafish larvae infected with *T. carassii* can be analysed at 4 dpi; based on up to 4 different parameters including 1) visual monitoring of larval behaviour, 2) parasite numbers, 3) location or 4) vasodilation, larvae could be segregated into high- and low-infected individuals. See details in the text in the corresponding Materials & Methods section.

### Real-time quantitative PCR

Zebrafish were sacrificed by an overdose of MS-222 anaesthetic (50 mg/L). At each time point 3-6 zebrafish larvae were sacrificed and pooled. Pools were transferred to RNA later (Ambion), kept at 4°C overnight and then transferred to −20°C for further storage. Total RNA isolation was performed with the Qiagen RNeasy Micro Kit (QIAgen, Venlo, The Netherlands) according to manufacturer’s protocol. Next, 250-500 ng total RNA was used as template for cDNA synthesis using SuperScript III Reverse Transcriptase and random hexamers (Invitrogen, Carlsbad, CA, USA), following the manufacturer’s instructions with an additional DNase step using DNase I Amplification Grade (Invitrogen, Carlsbad, CA, USA). cDNA was then diluted 25 times to serve as template for real-time quantitative PCR (RT-qPCR) using Rotor-Gene 6000 (Corbett Research, QIAgen), as previously described (Forlenza et al., 2012). Primers for *ef1a*: FW-CTGGAGGCCAGCTCAAACAT, RV-TCAAGAAGAGTAGTAGTACCG (ZDB-GENE-990415-52); *T. car. hsp70*: FW-CAGCCGGTGGAGCGCGT, RV-AGTTCCTTGCCGCCGAAGA (FJ970030.1, GeneBank) were obtained from Eurogentec (Liège, Belgium). Gene expression was normalized to the expression of *elongation factor-1 alpha (ef1a)* housekeeping gene and expressed relative to the PVP control at the same time point or to 0 days post injection (dpi) time point.

### *In vivo* imaging and videography of zebrafish

Prior to imaging, zebrafish larvae were anaesthetised with 0.017% MS-222 (Sigma-Aldrich). For total fluorescence acquisition, double transgenic *Tg(mpeg1.4:mCherry-F;mpx:GFP)* were positioned on preheated flat agar plates (1% agar in egg water with 0.017% MS-222) and imaged with Fluorescence Stereo Microscope (Leica M205 FA). The image acquisition settings were as following: Zoom: 2.0 - 2.2, Gain: 1, Exposure time (ms): 70 (BF)/700 (GFP)/1500 (mCherry), Intensity: 60 (BF)/700 (GFP)/700 (mCherry), Contrast: 255/255 (BF)/ 70/255 (GFP)/ 15/255 (mCherry).

Alternatively, anaesthetised larvae were embedded in UltraPure LMP Agarose (Invitrogen) and positioned on the coverglass of a 35 mm petri dish, (14 mm microwell, coverglass No. 0 (0.085-0.13mm), MatTek corporation) prior to imaging. A Roper Spinning Disk Confocal (Yokogawa) on Nikon Ti Eclipse microscope with 13×13 Photometrics Evolve camera (512 x 512 Pixels 16 x 16 micron) equipped with a 40x (1.30 NA, 0.24 mm WD) OI objective, was used with the following settings: GFP excitation: 491nm, emission: 496-560nm, digitizer: 200 MHz (12-bit); 561 BP excitation: 561nm; emission: 570-620nm, digitizer: 200 MHz (12-bit); BF: digitizer: 200 MHz (12-bit). Z-stacks of 1 or 0.5 μm. An Andor-Revolution Spinning Disk Confocal (Yokogawa) on a Nikon Ti Eclipse microscope with Andor iXon888 camera (1024 x 1024 Pixels 13 x 13 micron) equipped with 40x (0.75 NA, 0.66 mm WD) objective, 40x (1.15 NA, 0.61-0.59 mm WD) WI objective, 20x (0.75 NA, 1.0 mm WD) objective and 10x (0.50 NA, 16 mm WD) objective was used with the following settings: Dual pass 523/561: GFP excitation: 488nm, emission: 510-540nm, EM gain: 20-300ms, digitizer: 10 MHz (14-bit); RFP excitation: 561nm; emission: 589-628nm, EM gain: 20-300ms, digitizer: 10 MHz (14-bit); BF DIC EM gain: 20-300ms, digitizer: 10 MHz (14-bit). Z-stacks of 1 μm. Images were analysed with ImageJ-Fijii (version 1.52p).

High-speed videography of *T. carassii* swimming behaviour *in vivo* was performed as described previously (Dóró et al., 2019). Briefly, the high-speed camera was mounted on a DMi8 inverted digital microscope (Leica Microsystems), controlled by Leica LASX software (version 3.4.2.) and equipped with 40x (NA 0.6) and 20x (NA 0.4) long distance objectives (Leica Microsystems). For high-speed light microscopy a (8 bits) EoSens MC1362 (Mikrotron GmbH, resolution 1280 x 1024 pixels), with Leica HC 1x Microscope C-mount Camera Adapter, was used, controlled by XCAP-Std software (version 3.8, EPIX inc.). Images were acquired at a resolution of 900 x 900 or 640 x 640 pixels. Zebrafish larvae were anaesthetised with 0.017% MS-222 and embedded in UltraPure LMP Agarose (Invitrogen) on a microscope slide (1.4-1.6 mm) with a well depth of 0.5-0.8 mm (Electron Microscopy Sciences). Upon solidification of the agarose, the specimen was covered with 3-4 drops of egg water containing 0.017% MS-222, by a 24 x 50 mm coverslip and imaged immediately. For all high-speed videography, image series were acquired at 480–500 frames per second (fps) and analysed using a PFV software (version 3.2.8.2) or MiDAS Player v5.0.0.3 (Xcite, USA).

### Fluorescence quantification

Quantification of total cell fluorescence in zebrafish larvae was performed in ImageJ (version 1.49o) using the free-form selection tool and by accurately selecting the larvae area. Owing to the high auto-fluorescence of the gut or gut content, and large individual variation, the gut area was excluded from the total fluorescence signal. Area integrated intensity and mean grey values of each selected larva were measured by the software. To correct for the background, three consistent black areas were selected in each image. Analysis was performed using the following formula: corrected total cell fluorescence (CTCF) = Integrated density – (Area X Mean background value).

### EdU proliferation assay and immunohistochemistry

iCLICK™ EdU (5-ethynyl-2’-deoxyuridine, component A) from ANDY FLUOR 555 Imaging Kit (ABP Biosciences) at a stock concentration of 10 mM, was diluted in PVP to 1.13 mM. Infected *Tg(mpeg1:eGFP)* or *Tg(mpx:GFP)* larvae were injected in the heart cavity at 3 dpi (8dpf) with 2 nl of solution, separated in high- and low-infected individuals at 4 dpi and euthanized 6-8 hours later (30-32h after EdU injection) with an overdose of anaesthetic (0.4% MS-222 in egg water). Following fixation in 4% paraformaldehyde (PFA, Thermo Scientific) in PBS, o/n at 4°C, larvae were washed three times in buffer A (0.1% (v/v) tween-20, 0.05% (w/v) azide in PBS), followed by dehydration: 50% MeOH in PBS, 80% MeOH in H_2_0 and 100% MeOH, for 15 min each, at room temperature (RT), with gentle agitation. To remove background pigmentation, larvae were incubated in bleach solution (5% (v/v) H_2_O_2_ and 20% (v/v) DMSO in MeOH) for 1h at 4°C, followed by rehydration: 100% MeOH, 80% MeOH in H2O, 50% MeOH in PBS for 15 min each, at room temperature (RT), with gentle agitation. Next, larvae were incubated three times for 5 min each in buffer B (0.2%(v/v) triton-x100, 0.05% azide in PBS) at RT with gentle agitation followed by incubation in EdU iCLICK™ development solution for 30 min at RT in the dark and three rapid washes with buffer B.

The described EdU development led to loss of GFP signal in the transgenic zebrafish. Therefore, to retrieve the position of neutrophils or macrophages, wholemount immunohistochemistry was performed. Larvae were blocked in 0.2% triton-x100, 10% DMSO, 6% (v/v) normal goat serum and 0.05% azide in PBS, for 3h, at RT with gentle agitation. Next, the primary antibody Chicken anti-GFP (Aves labs.Inc., 1:500) in Antibody buffer (0.2% tween-20, 0.1% heparin, 10% DMSO, 3% normal goat serum and 0.05% azide in PBS) was added and incubated overnight (o/n) at 37°C. After three rapid and three 5 min washes in buffer C (0.1% tween-20, 0.1% (v/v) heparin in PBS), at RT with gentle agitation, the secondary antibody goat anti-chicken-Alexa 488 (Abcam, 1:500) was added in Antibody buffer and incubated o/n at 37°C. After three rapid and three 5 min washes in buffer C, at RT with gentle agitation, larvae were imaged with Andor Spinning Disk Confocal Microscope.

### BODIPY injection

BODIPY™ FL pentanoic acid (BODIPY-FL5, Invitrogen) was diluted in DMSO to a 3 mM stock solution. Stock solution was diluted 100x (30 μM) with PVP. Infected larvae 3 dpi (8 dpf) were injected with 1 nl of the solution i.p. (heart cavity) and imaged 18-20 hours later.

### Statistical analysis

Analysis of gene expression and total fluorescence data were performed in GraphPad PRISM 5. Statistical analysis of gene expression data was performed on Log(2) transformed values followed by One-way ANOVA and Dunnett’s multiple comparisons test. Analysis of Corrected Total Cell Fluorescence was performed on Log(10) transformed values followed by Two-way ANOVA and Bonferroni multiple comparisons post-hoc test. Analysis of EdU^+^ macrophages was performed on Log(10) transformed values followed by One-way ANOVA and Bonferroni multiple comparisons post-hoc test. In all cases, *p*<0.05 was considered significant.

## Results

### Susceptibility of zebrafish larvae to *T. carassii* infection

We recently reported the establishment of a trypanosome infection in zebrafish larvae using a natural fish parasite, *Trypanosoma carassii* (Dóró et al., 2019). To further investigate the immune response to *T. carassii* infection, we first investigated the kinetics of susceptibility of zebrafish larvae as well as the kinetics of expression of various immune-related genes. Similar to the previous report, *T. carassii* infection of 5 dpf zebrafish larvae leads to approximately 10-20% survival by 15 days post infection (dpi) with the highest incidence of mortality between 4 and 7dpi (**Fig 2A**). The onset of mortality coincided with the peak of parasitaemia as assessed by real-time quantitative gene expression analysis of a *T. carassii*-specific gene (**Fig 2B**). Nevertheless, we consistently observed 10-20% survival in the *T. carassii*-infected group, suggesting that zebrafish larvae can control *T. carassii* infection. This observation prompted us to investigate the kinetics of parasitaemia and development of (innate) immune responses at the individual level.

**Fig 2.**
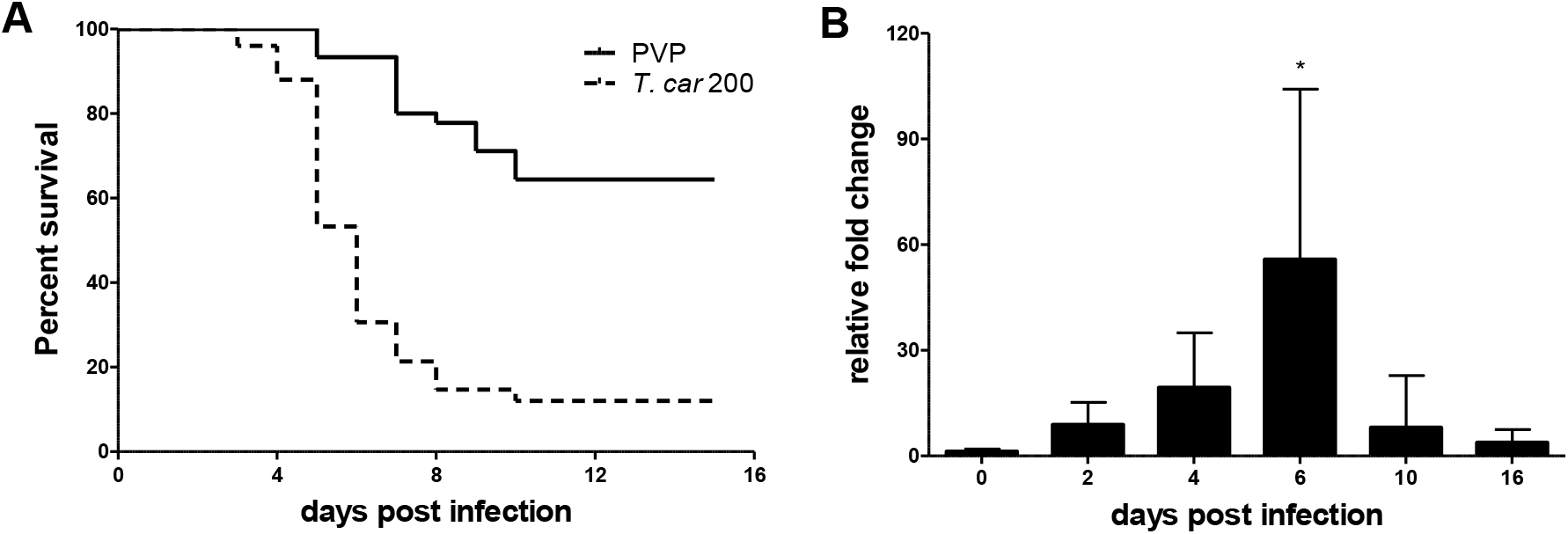
*T. carassii* infection of larval zebrafish. **A)** *Tg(mpeg1.4:mCherry-F;mpx:GFP)* larvae (5 dpf) were injected intravenously with *n*=200 *T. carassii/fish* or with PVP as control and survival was monitored over a period of 15 days. **B)** *Tg(mpeg1.4:mCherry-F;mpx:GFP)* zebrafish (5 dpf) were treated as in A and sampled at various time points. At each time point, 3-6 pools of 3-5 larvae were sampled for real-time quantitative PCR analysis. Relative fold change of the *T. carassii*-specific *heat-shock protein-70 (hsp70)* was normalised to the zebrafish-specific *ef1α* and expressed relative to the parasite injected group at time point 0h. Bars indicate average and standard deviation (SD) on n=3-6 pools per time point.

### Clinical signs of *T. carassii* infection and clinical scoring system

To characterize the response to *T. carassii* infection in individual zebrafish larvae, we developed a clinical scoring system to determine individual infection levels, enabling us to group individual larvae based on severity of infection. From 4 dpi onwards, we could consistently sort larvae into groups of high- or low-infected individuals based on *in vivo* observations, without the need to sacrifice animals (**S1 Video**). Infection levels were categorised using four criteria: 1) escape reflex (slow vs fast) upon contact with a pipette tip, 2) infection scores (1-10, see details in Materials and Methods), based on the ratio of blood cells and parasites passing through an intersegmental capillary (ISC) in 100 events (**Fig 3A, 3B**) (**S1 Video**, 00:06-00:39 sec), 3) extravasation, based on the presence of parasites outside of blood vessels (**Fig 3C**) (**S1 Video**, 00:40-1:20 sec) and 4) vasodilation, based on the diameter of the cardinal caudal vein (**Fig 3D, 3E**). The first criterion defined all individuals with a minimal escape reflex (slow swimmers) as high-infected individuals: they were mostly located at the bottom of the tank and showed minimal reaction upon direct contact with a pipette. Larvae with a normal escape reflex (fast swimmers) however, were not exclusively low-infected individuals. Therefore, a second criterion was used based on trypanosome counting in ISC (**S1 Video**, 00:06-00:39 sec). Individuals with an infection score between 1-3 were categorized as low-infected and always survived the infection (see Materials and Methods). Individuals with an infection score between 6-10 were categorized as high-infected and generally succumbed to the infection. Individuals with an intermediate score (4-5) could go both ways: they either showed a delayed parasitaemia and later developed high parasitaemia (common) or recovered from the infection (rare). The third criterion clearly identified high-infected individuals as those showing extensive extravasation at two or more of the following locations: peritoneal cavity (**Fig 3C**) (**S1 Video**, 00:40-00:59 sec), interstitial space lining the blood vessels, muscle tissue (**S1 Video**, 01:00-01:07 sec) or fins (**S1 Video**, 01:08-01:20 sec), in particular the anal fin. At these locations, in high-infected individuals, trypanosomes could accumulate in high numbers, filling up all available spaces. Extravasation however could also occur in low-infected individuals, but to a lesser extent. The fourth criterion, vasodilation of the cardinal caudal vein associated with high numbers of trypanosomes in the blood vessels, was a definitive sign of high infection level, and never occurred in low-infected larvae. To validate our scoring system, expression of a *T. carassii*-specific gene was analysed in pools of larvae classified as high- or low-infected. As expected, in individuals categorized as high-infected, *T. carassii*-specific gene expression increased more than 60-fold whereas in low-infected individuals the increase was less than 20-fold (**Fig 3F**). Altogether these data show that *T. carassii* infects zebrafish larvae, but that the infection can develop differently among individuals, leading to different outcomes. The clinical scoring system based on numerous criteria is suitable to reliably separate high- and low-infected larvae to further investigate individual immune responses.

**Fig 3.**
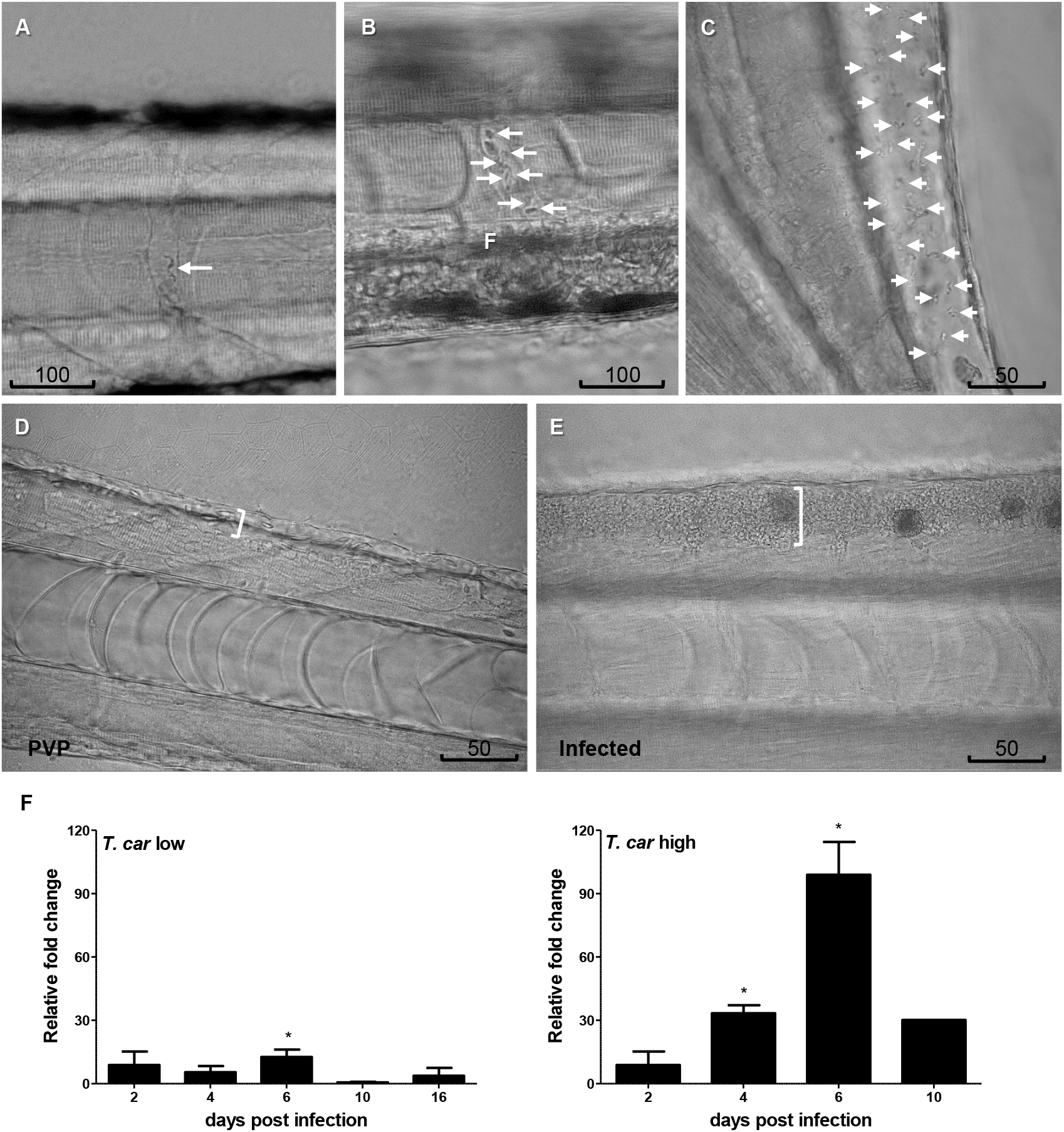
Progression of *T. carassii* infection in zebrafish larvae. *Tg(mpeg1.4:mCherry-F;mpx:GFP)* 5 dpf zebrafish were injected with *n*=200 *T. carassii* or with PVP and imaged at 2 dpi (**A**), 5 dpi (**B-C**), 7 dpi (**D-E**) or sampled at various time points after infection (**F**). Shown are representative images of intersegmental capillaries (ISC) containing various number of *T. carassii* (white arrows) (**A-B**); extravasated *T. carassii* (only some indicated with white arrows) in the intraperitoneal cavity (**C**); cardinal caudal vein diameter in PVP (**D**) or in *T. carassii*-infected larvae (**E**). Square brackets indicate the diameter of the cardinal caudal vein. Frames are extracted from high-speed videos acquired with a Leica DMi8 inverted microscope at a 40x magnification. **F)** *T. carassii* infection level. High- and low-infected individuals were separated from 4 dpi onwards based on our clinical scoring criteria. At each time point, 3 pools of 3-5 larvae were sampled for subsequent real-time quantitative gene expression analysis. Each data point represents the mean of 3 pools, except for the high-infected group at 10 dpi where only one pool could be made due to low survival. Relative fold change of the *hsp70* was normalised relative to the zebrafish-specific *ef1α* housekeeping gene and expressed relative to the trypanosome-injected group at time point 0h.

### *T. carassii* infection induces a strong macrophage response in zebrafish larvae

After having established a method to determine infection levels in each larva, we next investigated whether a differential innate immune response would be mounted in high- and low-infected fish. To this end, using double-transgenic *Tg(mpeg1.4:mCh-F;mpx:GFP)* zebrafish, we first analysed macrophage and neutrophil responses in whole larvae by quantifying total cell fluorescence in high- and low-infected individuals (**Fig 4**). Total neutrophil response (total green fluorescence) was not significantly affected by the infection (**Fig 4A, 4C**). In contrast, the macrophage response (total red fluorescence) increased significantly in infected individuals from 3 dpi onwards, and was most prominent in the head region and along the cardinal caudal vein (**Fig 4B**). In low-infected larvae, a significant increase in red fluorescence was observed already by 5 dpi and remained high up until 9 dpi; in high-infected larvae, despite a marginal but not significant increase at 5 and 7 dpi, significant differences were observed at day 9 after infection (**Fig 4C**). Interestingly, no significant differences were observed between high- or low-infected individuals, suggesting that despite the differences in trypanosome levels (**Fig 3F**), macrophages respond to the presence and not to the number of trypanosomes.

**Fig 4.**
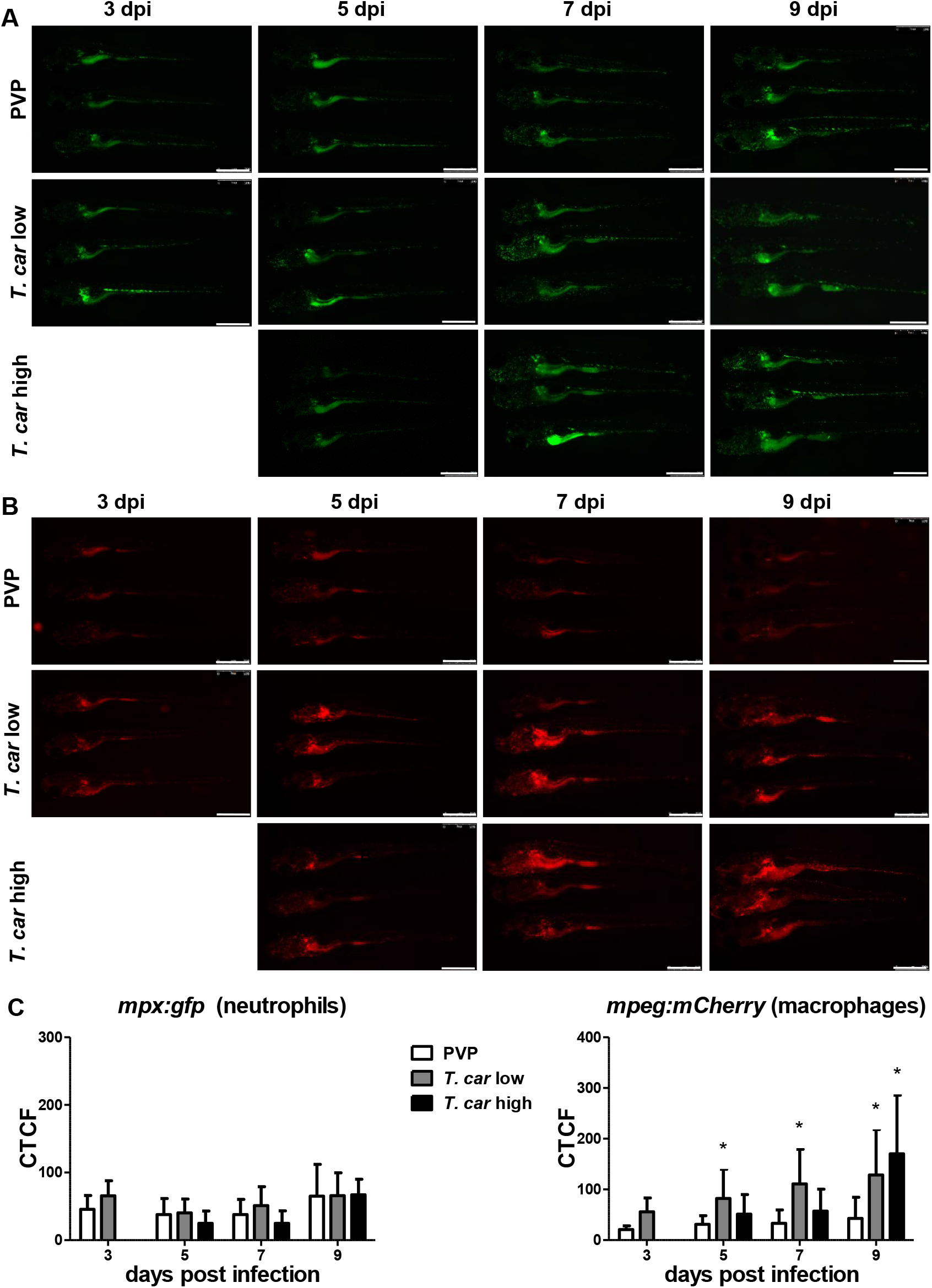
Macrophages respond more prominently than neutrophils to *T. carassii* infection. *Tg(mpeg1.4:mCherry-F;mpx:GFP)* were injected intravenously at 5 dpf with *n*=200 *T. carassii* or with PVP. At 4 dpi larvae were separated in high- and low-infected individuals. **A-B)** At the indicated time points, images were acquired with Leica M205FA Fluorescence Stereo Microscope with 1.79x zoom. Images are representatives of n=3-44 larvae per group, depending on the number of high- or low-infected larvae categorized at each time point. Scale bar indicates 750 μm. **C)** Corrected Total Cell Fluorescence (CTCF) quantification of infected and non-infected larvae. Owing to the high auto-fluorescence, the gut area was excluded from the total fluorescence signal as described in the methods section. Bars represent average and standard deviation of red and green fluorescence in n=5-44 whole larvae, from 2 independent experiments. * indicates significant differences (P<0.05) to the respective PVP control as assessed by Two-Way ANOVA followed by Bonferroni post-hoc test.

### *T. carassii* infection promotes macrophage and neutrophil proliferation

The increase in overall red fluorescence can be indicative of activation of the *mpeg* promotor driving mCherry expression, but also of macrophage proliferation. To address the latter hypothesis, *Tg(mpeg1:eGFP)* or *Tg(mpx:GFP)* zebrafish larvae were infected with *T. carassii*, and subsequently injected with iCLICK™ EdU for identification of proliferating cells. With respect to proliferation, developing larvae display a generalized high rate of cell division throughout the body that increases overtime particularly in hematopoietic organs such as the thymus or the head kidney. Thus, for a more sensitive quantification of the proliferative response of macrophages and neutrophils in response to the infection, EdU was injected at 3 dpi (8 dpf), and at 4 dpi, larvae were separated in high- and low-infected individuals, followed by fixation and whole mount immunohistochemistry 6-8h later (30-32h after EdU injection). This allowed evaluating the macrophage and neutrophil response right at the onset of the macrophage response observed in **Fig 4C** and concomitantly with the development of differences in parasitaemia. As expected, EdU^+^ nuclei could be identified throughout the body of developing larvae. When specifically looking at proliferating macrophage (**Fig 5**) and neutrophils (**Fig 6**) we selected the area of the head (left panels) and trunk (right panels) region, where previously (**Fig 4B**) the highest increase in red fluorescence was observed.

**Fig 5.**
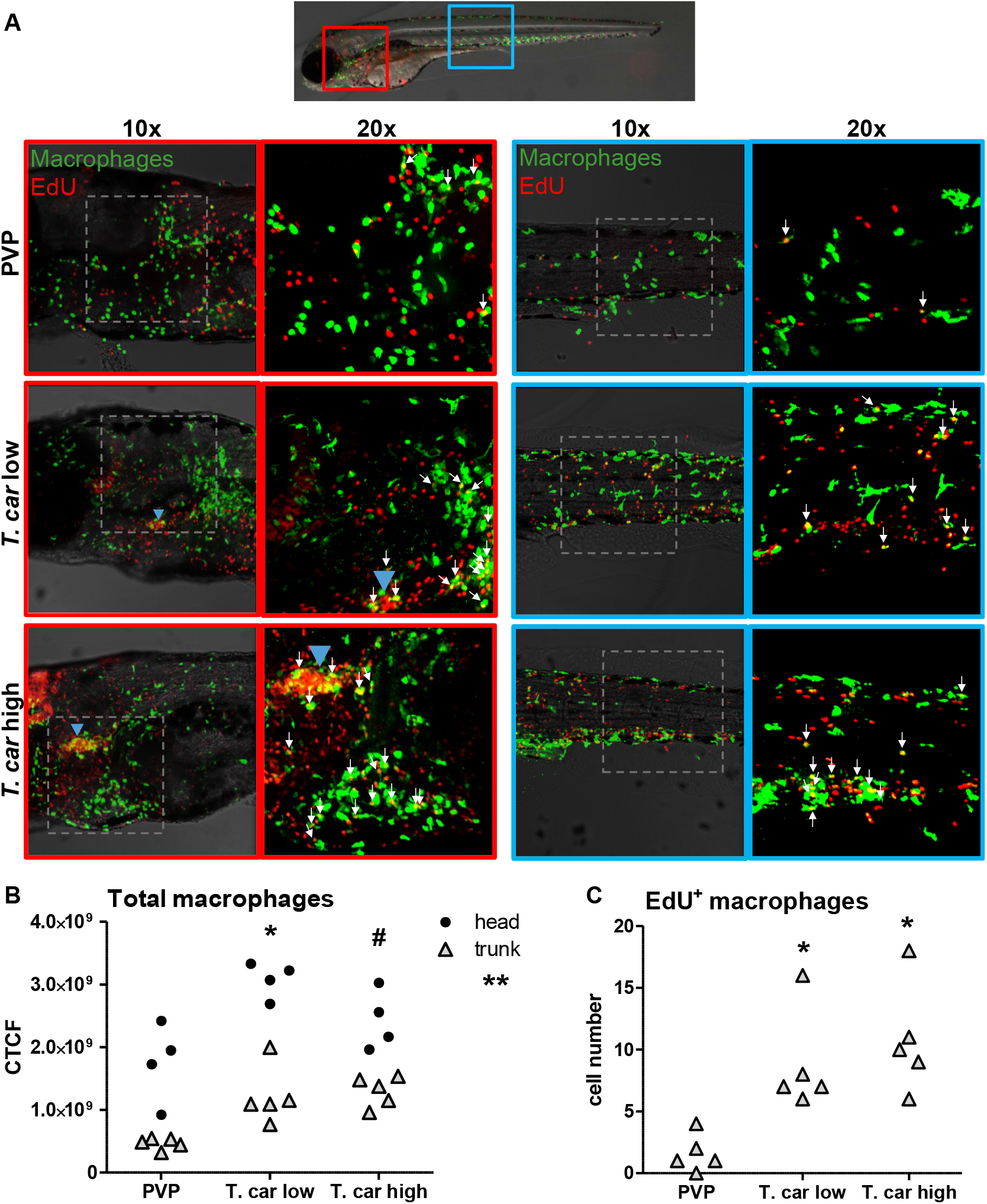
*T. carassii* infection triggers macrophage proliferation. *Tg(mpeg1:eGFP)* zebrafish larvae were infected intravenously at 5 dpf with n = 200 *T. carassii* or with PVP control (n=5 larvae per group, from four independent experiments). At 3 dpi, larvae received 2 nl 1.13mM iCLICK™ EdU, at 4 dpi were separated in high- and low-infected individuals and were imaged after fixation and whole mount immunohistochemistry 6-8h later (30-32h after EdU injection, ~9 dpf). Larvae were fixed and treated with iCLICK EdU ANDY FLUOR 555 (Red) development to identify EdU^+^ nuclei and with anti-GFP antibody to retrieve the position of macrophages, as described in the material and methods section. Larvae were imaged with Andor Spinning Disc Confocal Microscope using 10x and 20x magnification. **A)** Representative maximum projections of the head (left panels, red boxes) and trunk (right panels, blue boxes) regions capturing macrophages (green) and EdU^+^ nuclei (red) in PVP control, low- and high-infected zebrafish. In the PVP control group, EdU^+^ nuclei and GFP^+^ macrophages only rarely overlapped (white arrows, 20x), indicating limited proliferation of macrophages. In high- and low-infected individuals, the number of EdU^+^ macrophages increased (white arrows, 20x), indicating proliferation of macrophages in response to *T. carassii* infection. Blue arrowhead in the head of low and high-infected larvae, indicates the position of the thymus, an actively proliferating organ at this time point. The identification of EdU^+^ macrophages (white arrows) was performed upon detailed analysis of the separate stacks used to generate the overlay images, and are provided in **S2 Video**. **B)** Corrected total cell fluorescence (CTCF) calculated in the head (circles) and trunk (triangles) region. Symbols indicate individual larvae. * indicates significant differences of both, the head and trunk region to the respective PVP controls; # indicates significant differences of the trunk to the respective PVP control; ** indicates significant differences between CTCF in the head and trunk regions, as assessed by Two-Way ANOVA followed by Bonferroni post-hoc test. **C)** Number of EdU^+^ macrophages in the trunk region of infected and non-infected larvae. Symbols indicate individual larvae. * indicates significant differences to the PVP control as assessed by One-Way ANOVA followed by Bonferroni post-hoc test.

**Fig 6.**
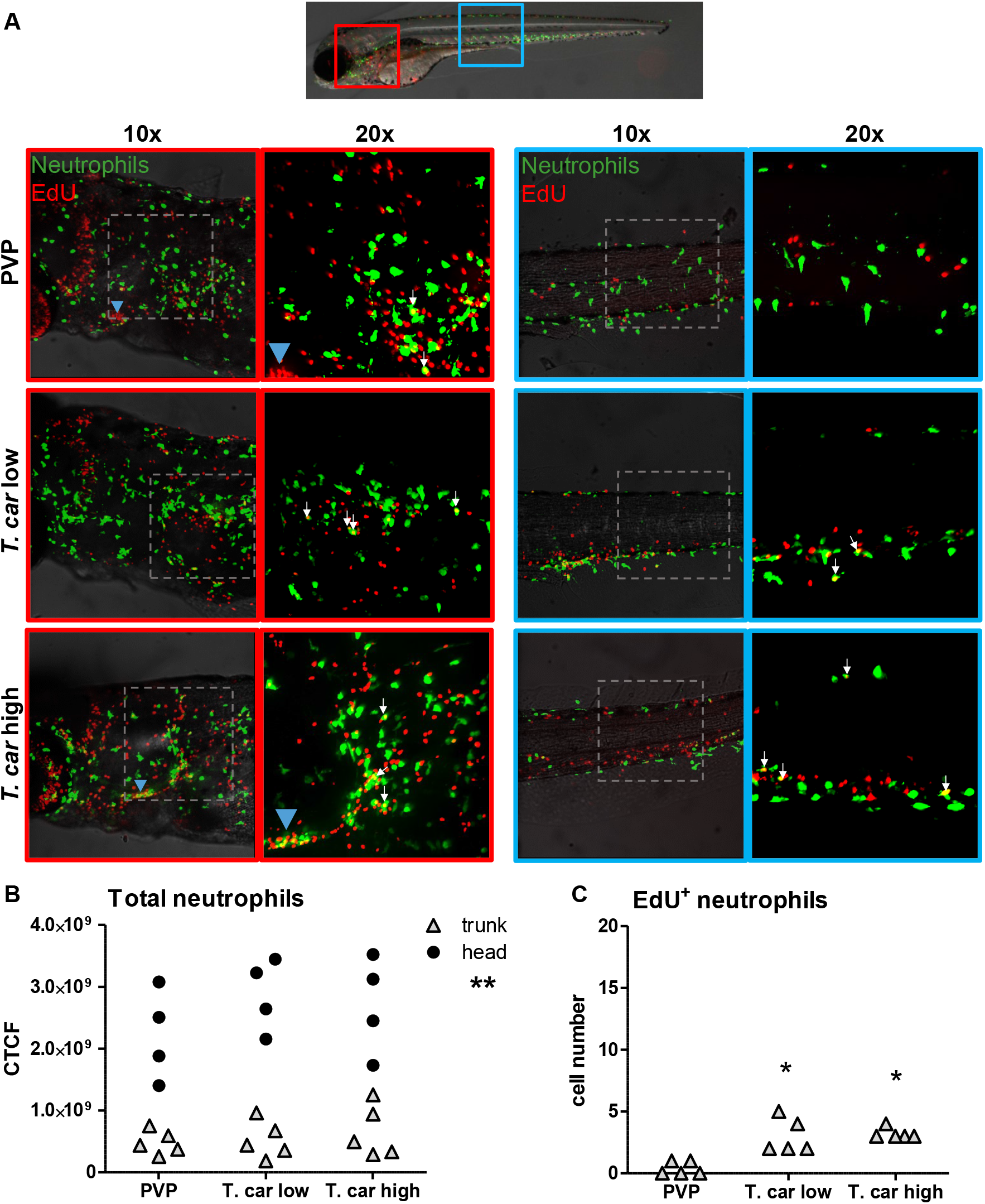
*T. carassii* infection triggers neutrophil proliferation. *Tg(mpx:GFP)* were treated as described in Fig 5 (n=5 larvae per group, from four independent experiments). **A)** Representative maximum projections of the head (left panels, red boxes) and trunk (right panels, blue boxes) region capturing neutrophils (green) and EdU^+^ nuclei (red) in PVP, low- and high-infected zebrafish. The images acquired at a 20x magnification show that in all groups, EdU^+^ nuclei and GFP^+^ neutrophils only rarely overlapped (white arrows), and was marginally higher in infected than in non-infected PVP controls. Detailed analysis of the separate stacks selected to compose the overlay image of the head region of the high-infected larva (bottom left panel), revealed that none of the neutrophils in the area indicated by the blue arrowhead (thymus) were EdU^+^ (**S3 Video**). **B)** Corrected total cell fluorescence (CTCF) calculated in the head (circles) and trunk (triangles) region. Symbols indicate individual larvae. ** indicates significant differences between CTCF in the head and trunk regions, as assessed by Two-Way ANOVA followed by Bonferroni post-hoc test. **C)** Number of EdU^+^ neutrophils in the trunk region of infected and non-infected larvae. Symbols indicate individual larvae. * indicates significant differences to the PVP control as assessed by One-Way ANOVA followed by Bonferroni post-hoc test.

When analysing the macrophage response, a greater number of macrophages was observed in the head and trunk of both high- and low-infected larvae compared to PVP-injected individuals (**Fig 5A**, 10x magnifications). In the head, macrophages were scattered throughout the region but in infected larvae they were most abundant in the area corresponding to the haematopoietic tissue (head kidney), posterior to the branchial arches, indicative of proliferation of progenitor cells. In the trunk, macrophages were scattered throughout the tissue, and in high-infected larvae in particular, macrophages also clustered in the cardinal caudal vein (**Fig 5A**, right panels). In agreement with previous observations (**Fig 4C**), quantification of total green fluorescence confirmed a significant increase in the head and trunk of low-infected larvae. In high-infected individuals, a significant increase was observed in the trunk, whereas in the head the number of macrophages was clearly elevated although not significantly when compared to the PVP-injected controls (**Fig 5B**). In all groups, total cell fluorescence in the head region was significantly higher than in the trunk region (**Fig 5B**), and thus largely contributed to the total cell fluorescence previously measured in whole larvae (**Fig 4C**). The difference in CTCF values between Fig 4 and Fig 5 can be attributed to the different microscopes and magnification used for the acquisition as well as fluorescence source (GFP or mCherry in Fig 4 and Alexa-488 fluorophore in Fig 5).

Given the high number of macrophages in the head region, their heterogeneous morphology, the thickness of the tissue and the overall high number of EdU^+^ nuclei, it was not possible to reliably count single (EdU^+^) macrophages in this area. Therefore, when analysing the degree of proliferation, we focused on the trunk region only. There, EdU^+^ macrophages could be observed in all groups, and in agreement with the total cell fluorescence measured in the same region (**Fig 5B**), their number was higher in low- and high-infected individuals compared to PVP-injected controls (**Fig 5C** and corresponding **S2 Video)**. No significant difference was observed between high- and low-infected fish, confirming that macrophages react to the presence and not to the number of trypanosomes. Within the trunk region of high-infected larvae, a large proportion of macrophages were observed around and inside the cardinal caudal vein, the majority of which were EdU^+^ (**S1A Fig**), suggesting that in high-infected larvae, proliferating macrophages migrated to the vessels. Altogether, these data confirm that *T. carassii* infection triggers macrophage proliferation and that proliferation is higher in low-infected compared to high-infected individuals, possibly due to a higher haematopoietic activity. When analysing the neutrophils response, in agreement with the previous observation, the number of neutrophils in the head and trunk regions was not apparently different between infected and non-infected larvae (**Fig 6A**). Neutrophils were scattered throughout the head region, but differently from macrophages, their number did not increase in the area corresponding to the haematopoietic tissue. Quantification of total cell fluorescence in the head and trunk revealed no significant differences between groups (**Fig 6B, S3 Video)**. Interestingly, quantification of EdU^+^ neutrophils in the trunk region, revealed that while in PVP-injected individuals EdU^+^ neutrophils were rarely observed, in infected fish, a significant, although low number of EdU^+^ neutrophils was present (**Fig 6C**). These data indicate that neutrophils also respond to the infection by proliferating, but their number is relatively low and may have not significantly contributed to changes in total cell fluorescence. In contrast to macrophages, within the analysed trunk region, neutrophils were never observed within the cardinal caudal vein, and independently of whether they proliferated (EdU^+^) or not, were mostly observed lining the vessel (**S1B Fig**). Altogether, these data indicate that independent of the trypanosome number, *T. carassii* triggers a differential macrophage and neutrophil proliferation, where macrophages respond more prominently than neutrophils to the infection.

### Differential distribution of neutrophils and macrophages in high- and low-infected zebrafish larvae

After having established that *T. carassii* infection triggers macrophage, and to a lesser extent, neutrophil proliferation, we next investigated whether a differential distribution of these cells occurred during infection. Considering that trypanosomes are blood dwelling parasites and the kinetics of parasitaemia, we focused on the cardinal caudal vein at 4 dpi, a time point at which clear differences in parasitaemia (**Fig 3**) and a differential distribution of macrophages and neutrophils (**Fig 5–6** and **S1 Fig**) were observed between high- and low-infected larvae. To this end, crosses between transgenic lines marking the blood vessels and those marking either macrophages or neutrophils were used. *Tg(kdrl:caax-mCherry;mpx:GFP)* or *Tg(fli1:eGFP x mpeg1.4:mCherry-F)* were infected with *T. carassii*, separated into high- and low-infected larvae at 4 dpi, and imaged with Roper Spinning Disk Confocal Microscope using 40x magnification. Longitudinal and orthogonal images of the vessel were analysed to visualise the exact location of cells along the cardinal caudal vein (**Fig 7** and **S4 Video**). In PVP controls, both macrophages and neutrophils were exclusively located outside the vessel in close contact with the endothelium or in the tissue adjacent the cardinal caudal vein. In infected fish, while neutrophils remained exclusively outside the vessel (**Fig 7**, left panel), macrophages could be seen both inside (white arrows) and outside (blue arrows) the vessel (**Fig 7**, right panel). Whilst in low-infected individuals macrophage morphology was similar to that observed in non-infected fish, in high-infected larvae, macrophages inside the vessel clearly had a more rounded morphology. Altogether these data indicate that differently from neutrophils, macrophages increase in number in infected fish, are recruited inside the cardinal caudal vein and, depending on the infection level, their morphology can be greatly affected.

**Fig 7.**
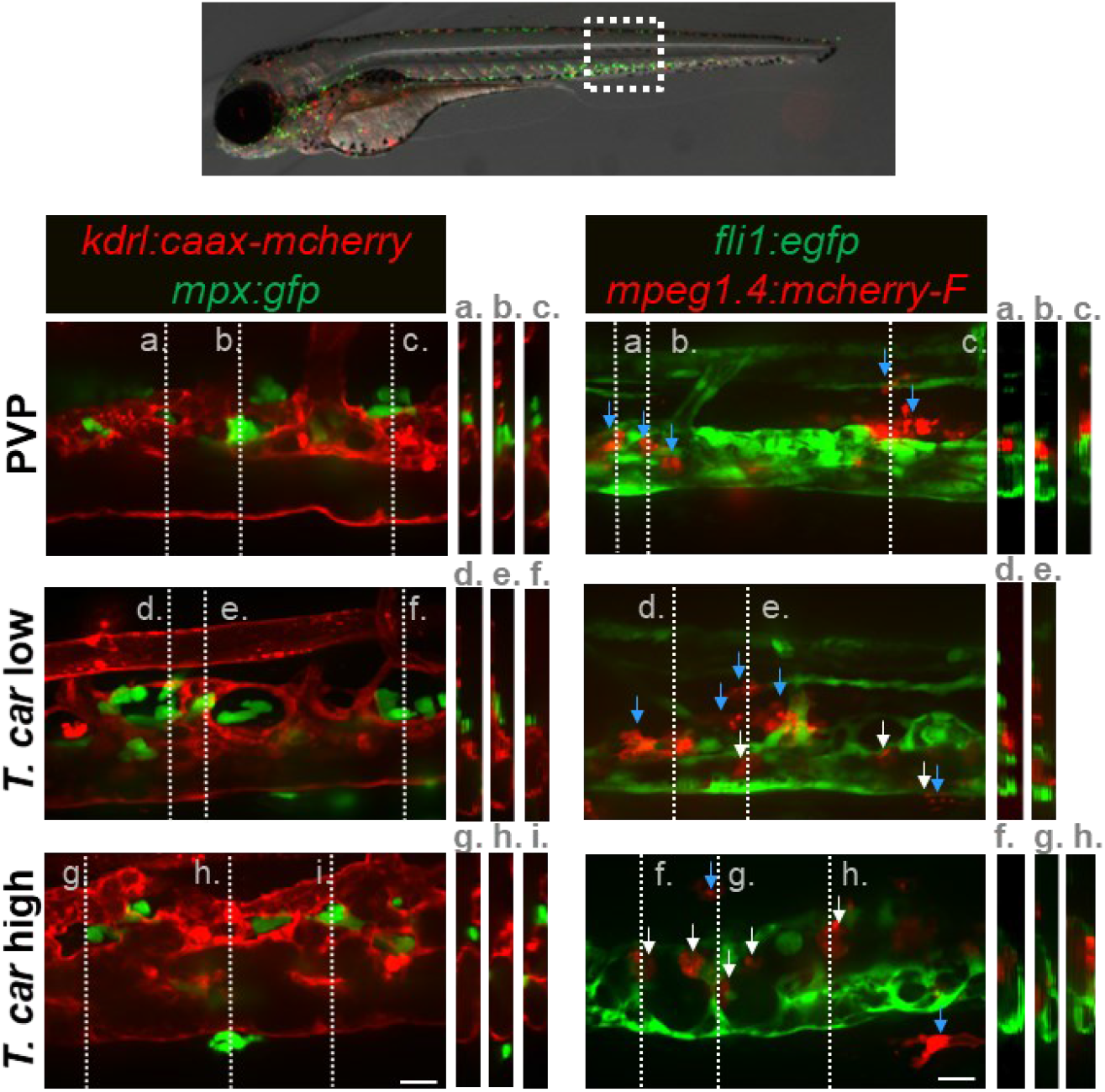
Macrophages are recruited into the cardinal caudal vein of high-infected zebrafish larvae. *Tg(kdrl:caax-mCherry;mpx:GFP)* and *Tg(fli1:eGFP x mpeg1.4: mCherry-F)* zebrafish larvae (n=5 larvae per group, from two independent experiments) were injected intravenously at 5 dpf with n=200 *T. carassii* or with PVP. At 4 dpi larvae were separated in high- and low-infected groups and imaged with a Roper Spinning Disk Confocal Microscope using 40x magnification. Scale bars indicate 25 μm. **Left panel**: representative images of the longitudinal view of the cardinal caudal vein (red), capturing the location of neutrophils (green). Orthogonal views of the locations marked with grey dashed lines (a,b,c,d,e,f,g,h,i), confirm that in all groups, neutrophils are present exclusively outside the vessel. **Right panel**: representative images of the longitudinal view on the vessel, capturing the position of macrophages (red) outside (blue arrowheads) or inside (white arrowheads) the vessel (green). Orthogonal views of the locations marked with grey dashed lines (a,b,c,d,e,f,g,h) confirm that in PVP controls, macrophages are present exclusively outside the vessel (blue arrows); in low-infected larvae, most macrophages are outside the vessel (blue arrows) having an elongated or dendritic morphology, although seldomly rounded macrophages can be observed within the vessel (white arrows); in high-infected larvae, although macrophages with dendritic morphology can be seen outside the vessel, the majority of the macrophages resides inside the vessel, clearly having a rounded morphology. **S4 Video** provides the stacks used for the orthogonal views.

### *T. carassii* infection triggers the formation of foamy macrophages in high-infected fish

When analysing macrophage morphology and location, clear differences could be observed between control and high- or low-infected larvae when examined in greater detail. In control fish, macrophages generally exhibited an elongated and dendritic morphology, were very rarely observed inside the vessel and were mostly located along the vessel endothelium, in the tissue lining the cardinal vessels or in the ventral fin (**Fig 8A**, left). A similar morphology and distribution were observed in low-infected larvae (not shown). Strikingly, in high-infected larvae, we consistently observed large, dark, granular and round macrophages located almost exclusively inside the cardinal vein on the dorsal luminal side. These dark macrophages were clearly visible already in bright field images due to their size, colour and location, and could be present as single cells or as aggregates (**Fig 8A**, right). The occurrence of these large macrophages increased with the progression of the infection and were exclusive to high-infected individuals (**S5 Video**) as were never observed in low-infected or control larvae.

**Fig 8.**
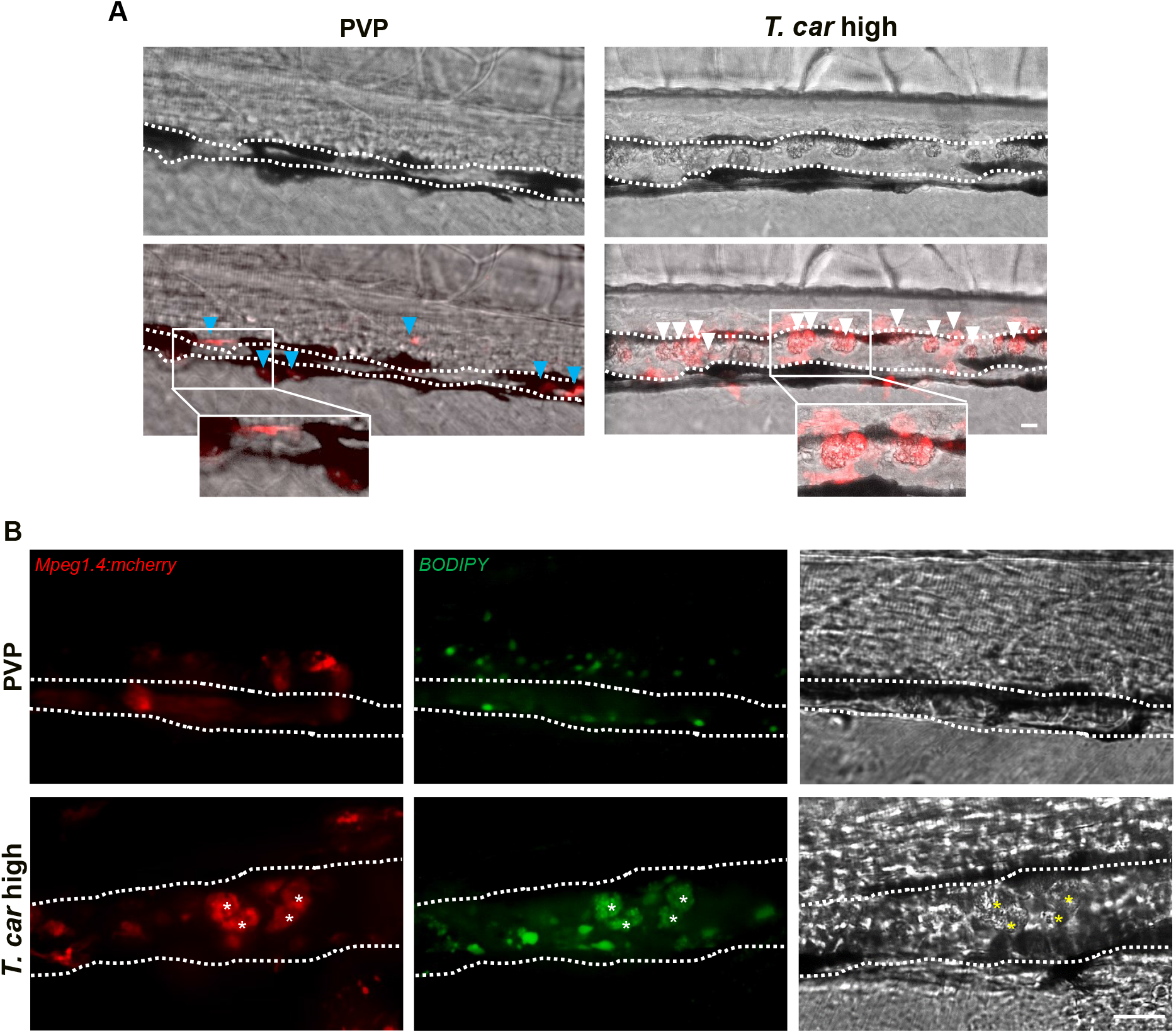
The large macrophages inside the cardinal caudal vein of high-infected zebrafish are foamy macrophages. **A)** *Tg(mpeg1.4:mCherry-F; mpx:GFP)* zebrafish larvae were infected intravenously at 5 dpf with *n*=200 *T. carassii* or with PVP (n=5 larvae per group) and imaged at 7 dpi using an Andor Spinning Disc Confocal Microscope at a 20x magnification. Representative images from three independent experiments are shown, with blue arrowheads pointing at macrophages outside the vessel and white arrowheads indicating large round macrophages inside the cardinal vein (dashed line). Scale bar indicates 25 μm. **B)** *Tg(mpeg1.4:mCherry-F)* were treated as in A (n=5 larvae per group). At 3 dpi, larvae received 1 nl of 30 μM BODIPY-FLC5 and were imaged 18-20 hours later using a Roper Spinning Disc Confocal Microscope at a 40x magnification. Representative images from three independent experiments are shown. Scale bar indicates 25 μm. **S6 Video** provides the stacks used in B.

The rounded morphology, size and dark appearance of these cells was reminiscent of that of foamy macrophages. Therefore, to further investigate the nature of these cells, the green fluorescent fatty acid BODIPY-FLC5 was used to track lipid accumulation in infected larvae (**Fig 8B**). Interestingly, administration of BODIPY-FL5 in high-infected larvae, one day prior to the expected appearance of the large macrophages, revealed the accumulation of lipids in these cells (**S6 Video**). Macrophages without this large, dark, granular appearance did not show lipid accumulation. This result therefore confirms that the large, rounded, granular macrophages in the cardinal caudal vein are indeed foamy macrophages.

### Foamy macrophages have a pro-inflammatory activation state

To further investigate the activation state of foamy macrophages, we made use of the *Tg(tnfa:eGFP-F;mpeg1.4:mCherry-F)* and *Tg(il1b:eGFP-F x mpeg1.4:mCherry-F)* double transgenic zebrafish lines, having macrophages in red and *tnfa*- or *Il1b*-expressing cells in green (**Fig 9** and **Fig 10**). We first focused on the time point at which the foamy macrophages were most clearly present in highly infected individuals, 4 dpi. Our results clearly show that all large foamy macrophages, were strongly positive for *tnfa*, suggesting an inflammatory activation state (**Fig 9A**). Interestingly, not only the large foamy macrophages within the vessel, but also dendritic or lobulated macrophages outside or lining the vessel showed various degrees of activation. Macrophages that were still partly in the vessel (**Fig 9B**, yellow arrowhead) displayed higher *tnfa* expression than macrophages lining the outer endothelium (white arrow heads). This could suggest that the presence of *T. carassii* components within the vessels might trigger macrophage activation.

**Fig 9.**
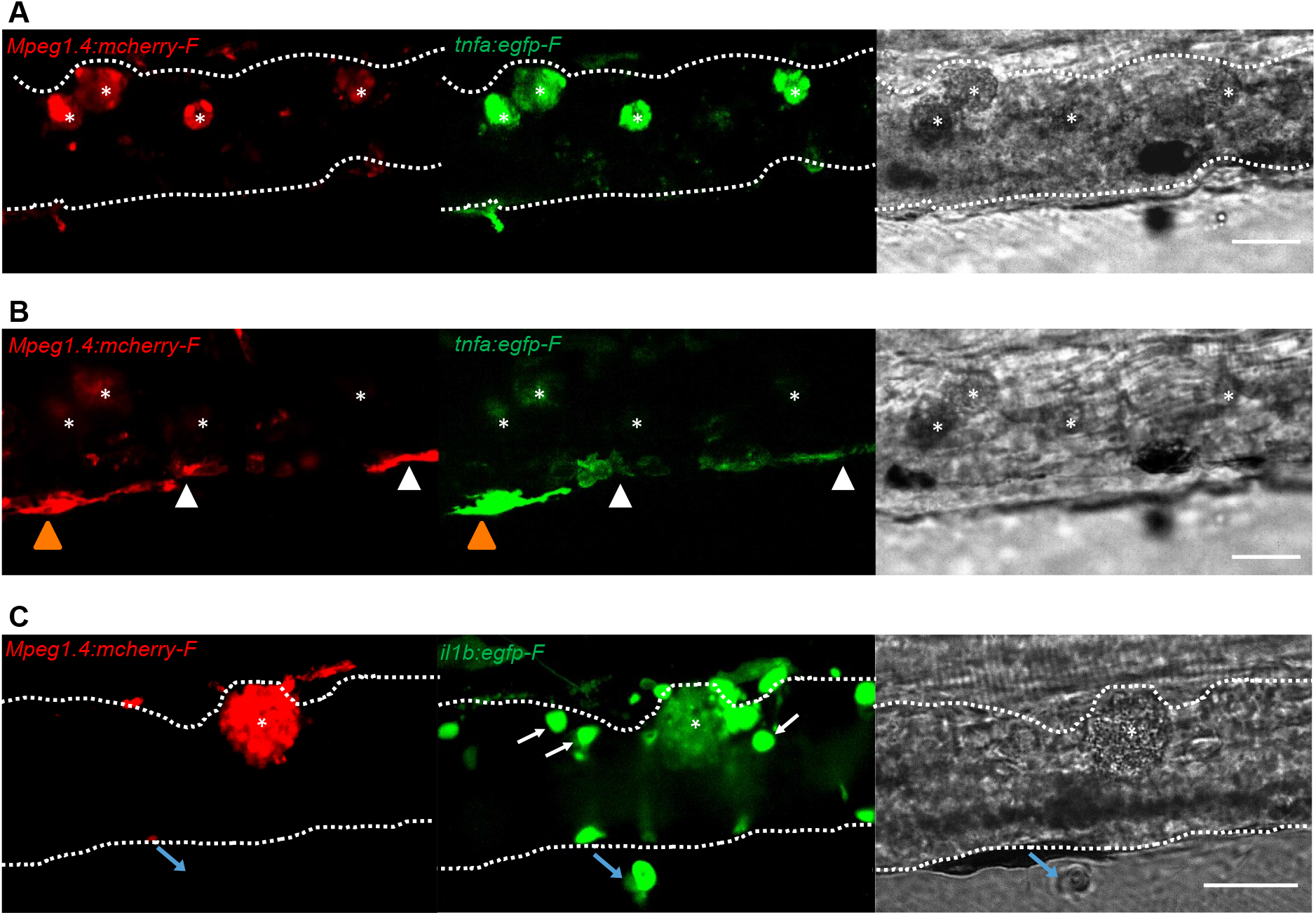
Foamy macrophages have an inflammatory profile. *Tg(tnfa:eGFP-F;mpeg1.4:mCherry-F)* **A-B**) or *Tg(il1b:eGFP-F x mpeg1.4:mcherry-F)* **C**) zebrafish larvae (5dpf), were injected with n = 200 *T. carassii* or with PVP. At 4 dpi, high-infected individuals were imaged with an Andor (**A-B**) or Roper (**C**) Spinning Disk Confocal Microscope using 40x magnification. Scale bar indicates 25 μm. Foamy macrophages (asterisks) were easily identified within the cardinal caudal vessel (dashed lines) and were strongly positive for *tnfa* (**A**) and *il1b* (**C**) expression (GFP signal). **B**) Same as in A, but a few stacks up, focusing on the cells lining the endothelium. Macrophages that were partly inside and partly outside the vessel (yellow arrowhead) were also strongly positive for *tnfa*, whereas macrophages lining the outer endothelium had a lower *tnfa* expression (white arrowheads). **C**) A foamy macrophage (asterisk) within the cardinal caudal vessel (dashed lines) was positive for *il1b*. Endothelial cells were also strongly positive for *il1b*, a selection of which is indicated by white arrows. A mCherry-negative-Illb positive cell is present outside the vessel (blue arrow). Given its position, it is likely to be a neutrophil.

**Fig 10.**
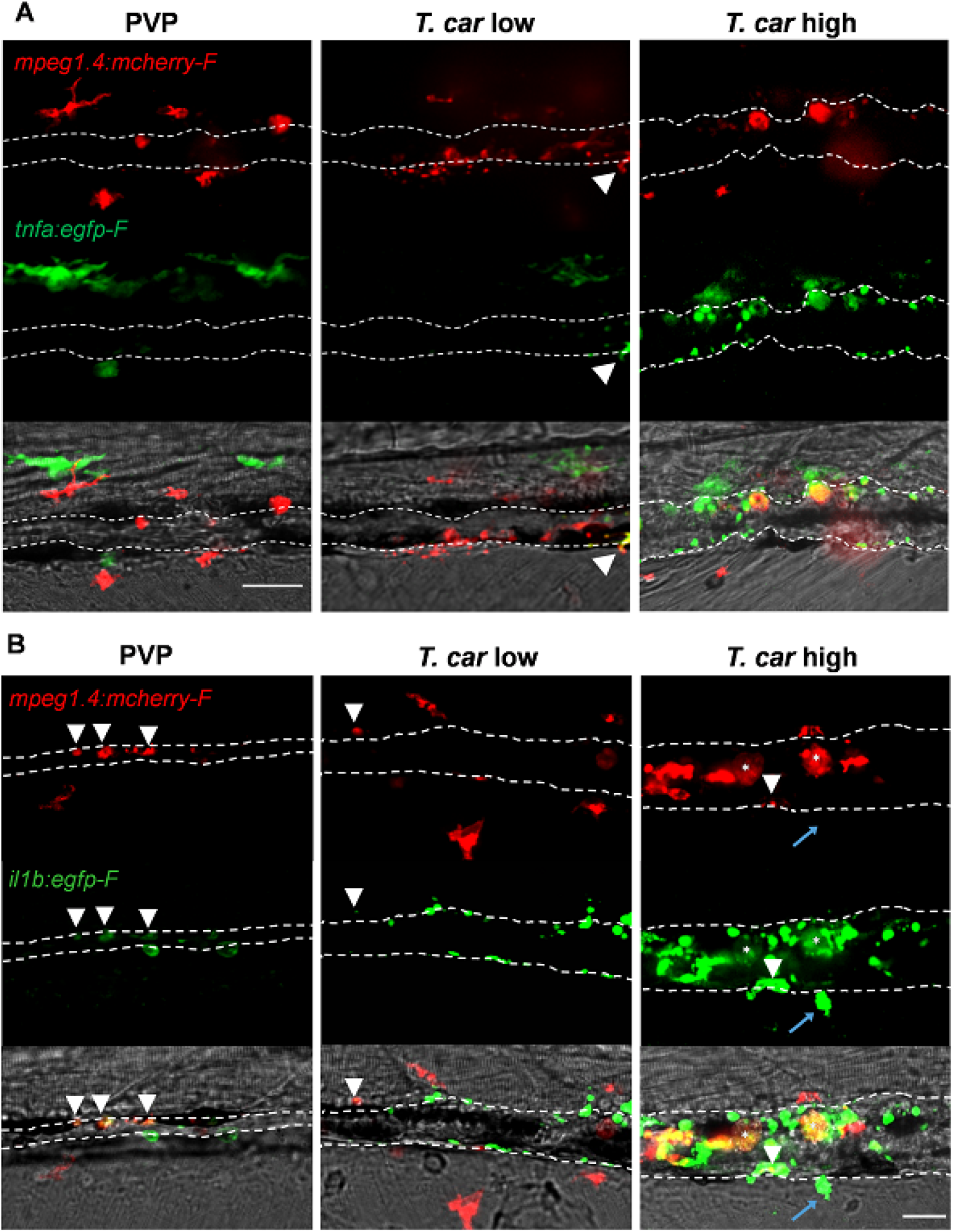
High-infected zebrafish have a strong inflammatory profile. Zebrafish larvae (5 dpf), either (**A**) *Tg(tnfa:eGFP-F x mpeg1.4:mCherry-F)* (n= 8-13 larvae per group from four independent experiments), or (**B**) *Tg(il1b:eGFP-F; mpeg1.4:mCherry-F)* (n=7-8 larvae per group from two independent experiments), were infected as described in Fig 7. At 3 dpi, larvae were separated in high- and low-infected individuals and at 4 dpi imaged with a Roper Spinning Disk Confocal Microscope. Scale bar indicate 25 μm. **A)** In non-infected PVP controls (left panel), several macrophages can be observed outside the vessel but none was positive for *tnfa*. In low-infected individuals (middle panel) macrophages were present inside and outside the vessel. Except the occasional macrophage showing *tnfa*-egfp expression (white arrowhead), they generally did not exhibit strong GFP signal. In high-infected individuals however, foamy macrophages (asterisks) as well as endothelial cells (bright green cells) or other leukocytes, were strongly positive for *tnfa*-egfp expression. **B)** *il1b*-*egfp* expression was generally low in non-infected PVP controls. In low-infected larvae *il1b* positive macrophages were rarely observed (white arrowhead). In both high- and low-infected fish, some endothelium cells in the cardinal caudal vein show high *il1b*-egfp expression (bright green cells in middle and right panel). In high-infected individual however (right panel), foamy macrophages inside the vessel (asterisks) as well as other macrophages lining the vessel (white arrowhead) and leukocytes in the tissue (blue arrow), were positive for *il1b*-egfp expression.

Similar to what observed for *tnfa* expression, all foamy macrophages within the vessel were also positive for *il1b* (**Fig 9C**, asterisk), confirming their pro-inflammatory profile. Interestingly, not only macrophages but also endothelial cells (a selection is indicated by white arrows) were strongly positive for *il1b*. Outside the vessel, cells that were mCherry negative but strongly positive for *Il1b* could also be observed (**Fig 9C**, blue arrow); given their position outside the vessel, these are most likely neutrophils. Altogether these data indicate that foamy macrophages occur in high-infected larvae and have a strong pro-inflammatory profile.

### High-infected zebrafish have a strong inflammatory profile associated with susceptibility to infection

When comparing the overall inflammatory state in high- and low-infected larvae it was apparent that high-infected individuals generally exhibited a higher pro-inflammatory response (**Fig 10**). Although *tnfa*- and *il1b-positive* macrophages could be seen in low-infected individuals, these were generally few (**Fig 10A** and **10B**, middle panels) and a higher number of *tnfa*- and *il1b*-expressing cells was observed in high-infected larvae (**Fig 10A** and **10B**, right panels). In these fish, *il1b* and *tnfa* expression was observed not only in (foamy) macrophages (asterisk), but also in mCherry negative cells outside the vessel (blue arrow, likely neutrophils) and in endothelial cells lining the vessel (bright green). As mentioned earlier, high-infected individuals are not able to control parasitaemia and generally succumbed to the infection. Altogether, these results suggest that in high-infected individuals, uncontrolled parasitaemia leads to an exacerbated pro-inflammatory response leading to susceptibility to the infection. Low-infected individuals however, with increased macrophage number and moderate cytokine responses, are able to control parasitaemia and to recover from the infection.

## Discussion

In this study we describe the differential response of macrophages and neutrophils *in vivo*, during the early phase of trypanosome infection of larval zebrafish. Considering the prominent role of innate immune factors in determining the balance between pathology and control of first-peak parasitaemia in mammalian models of trypanosomiasis (Magez and Caljon, 2011; Radwanska et al., 2018; Stijlemans et al., 2017), the use of transparent zebrafish larvae, devoid of a fully developed adaptive immune system, allowed us to investigate the early events of the innate immune response to *T. carassii* infection *in vivo*. After having established a clinical scoring system of infected larvae, we were able to consistently differentiate high- and low-infected individuals, each associated with opposing susceptibility to the infection. In high-infected larvae, which fail to control first-peak parasitaemia, we observed a strong inflammatory response associated with the occurrence of foamy macrophages and susceptibility to the infection. Conversely, in low-infected individuals, which succeeded in controlling parasitaemia, we observed a moderate inflammatory response associated with resistance to the infection. Altogether these data confirm that also during trypanosome infection of zebrafish, innate immunity is sufficient to control first-peak parasitaemia and that a controlled inflammatory response is beneficial to the host.

Using transgenic lines marking macrophages and neutrophils, total cell fluorescence and cell proliferation analysis revealed that *T. carassii* infection triggers macrophage proliferation, particularly in low-infected individuals. Although to a much lesser extent, neutrophils also responded to the infection by proliferating. The total number of neutrophils however, was comparatively low and likely did not contribute to the total cell fluorescence measured in our whole larvae analysis. Although neutrophils were recently implicated in promoting the onset of tsetse fly-mediated trypanosome infections in mouse dermis, macrophage-derived immune mediators, such as NO and TNFα were confirmed to played a more prominent role in the control of first-peak parasitaemia and in the regulation of the overall inflammatory response (Caljon et al., 2018).

The observation that in low-infected individuals the number of macrophages was significantly increased by 4-5 dpi, the time point at which clear differences in parasitaemia were apparent between the two infected groups, suggests a role for macrophages, or for macrophage-derived factors in first-peak parasitaemia control. Phagocytosis however, can be excluded as one of the possible contributing factors since motile *T. carassii*, similar to other extracellular trypanosomes (Caljon et al., 2018; Saeij et al., 2003; Scharsack et al., 2003), cannot be engulfed by any innate immune cell. A strong inflammatory response is also not required for trypanosomes control, since in low-infected individuals, only moderate *il1b* or *tnfα* expression was observed, mostly in macrophages, as assessed using transgenic zebrafish reporter lines. Our data are in agreement with several previous studies using trypanoresistant (BALB/c) or trypanosusceptible (C57Bl/6) mice that revealed the double-edge sword of pro-inflammatory mediators such as TNFα or IFNγ during trypanosome infection in mammalian models (reviewed by Radwanska et al., 2018; Stijlemans et al., 2007). These studies showed that a timely but controlled expression of IFNγ, TNFα and NO, contributed to trypanosomes control via direct (Daulouede et al., 2001; F Iraqi et al., 2001; Lucas et al., 1994) or indirect mechanisms (Kaushik et al., 1999; Magez et al., 2007, 2006; Mansfield and Paulnock, 2005; Namangala et al., 2001; Noël et al., 2002). Conversely, in individuals in which an uncontrolled inflammatory response took place, immunosuppression and inflammation-related pathology occurred (Namangala et al., 2009, 2001; Noël et al., 2004; Stijlemans et al., 2016). The stark contrast between the mild inflammatory response observed in low-infected individuals and the exacerbated response observed in high-infected larvae, strongly resembles the opposing responses generally observed in trypanoresistant or trypanosusceptible animal models. Owing to the possibility to monitor the infection at the individual level, it was possible to observe such responses within one population of outbred zebrafish larvae. Although we were unable to investigate the specific role of Tnfα during *T. carassii* infection of zebrafish, due to the unavailability of *tnfα-/-* zebrafish lines or the unsuitability of morpholinos for transient knock-down at late stages of development, we previously reported that recombinant zebrafish (as well as carp and trout) Tnfα, are all able to directly lyse *T. brucei* (Forlenza et al., 2009). In the same study, we reported that also during *Trypanoplasma borreli* (kinetoplastid) infection of common carp, soluble as well as transmembrane carp Tnfα play a crucial role in both trypanosome control and susceptibility to the infection. Thus, considering the evolutionary conservation of the lectin-like activity among vertebrate’s TNFα (Daulouede et al., 2001; Forlenza et al., 2009; R Lucas et al., 1994; Magez et al., 1997) it is possible that the direct lytic activity of zebrafish Tnfα may have played a role in the control of first-peak parasitaemia in low-infected individuals. In the future, using *tnfα*-/- zebrafish lines, possibly in combination with *ifnγ* reporter or *ifnγ-/-* lines, it will be possible to investigate in detail the relative contribution of these inflammatory mediators in the control of parasitaemia as well as onset of inflammation.

There are multiple potential explanations for the inability of high-infected larvae to control parasitaemia and the overt inflammatory response. Using various comparative mice infection models, it became apparent that while TNFα production is required for parasitaemia control, a timely shedding of TNFα Receptor-2 (TNFR2) is necessary to limit TNFα-mediated infection-associated immunopathology (Radwanska et al., 2018). Furthermore, during *T. brucei* infection in mice and cattle, continuous cleavage of membrane Glycosyl-phosphatidyl-inisotol (GPI)-anchored VSG (mVSG-GPI) leads to shedding of the soluble VSG-GIP (sVSG-GPI), while the di-myristoyl-glycerol compound (DMG) is left in the membrane. While the galactose-residues of sVSG-GPI constituted the minimal moiety required for optimal TNFα production, the DMG compound of mVSG contributed to macrophage overactivation (TNFα and IL-1β secretion) (Magez et al., 2002, 1998; Sileghem et al., 2001). Although *T. carassii* was shown to possess a surface dominated by GPI-anchored carbohydrate-rich mucin-like glycoproteins, not subject to antigenic variation (LISCHKE et al., 2000; Overath et al., 2001), components of its excreted/secreted proteome, together with phospholipase C-cleaved GPI-anchored surface proteins, have all been shown to play a role in immunogenicity (Joerink et al., 2007), inflammation (Oladiran and Belosevic, 2010, 2009; Ribeiro et al., 2010) as well as immunosuppression (Oladiran and Belosevic, 2012). Thus, the overactivation caused by the presence of elevated levels of pro-inflammatory trypanosome-derived moieties, combined with the lack of a timely secretion of regulatory molecules (e.g. soluble TNFR2) that could control the host response, may have all contributed to the exacerbated inflammation observed in high-infected individuals.

Given the differential response observed in low- and high-infected individuals, especially with respect to macrophage distribution and activation, we attempted to investigate the specific role of macrophages in the protection or susceptibility to *T. carassii* infection. To this end, the use of a cross between the *Tg(mpeg1:Gal4FF)^gl2^* (Ellett et al., 2011) and the *Tg(UAS-E1b:Eco.NfsB-mCherry)^c26^* (Davison et al., 2007) line, which would have allowed the timed metronidazole (MTZ)-mediated depletion of macrophages in zebrafish larvae, was considered. Unfortunately, *in vitro* analysis of the effect of MTZ on the trypanosome itself, revealed that trypanosomes are susceptible to MTZ, rendering the *nfsB* line not suitable to investigate the role of macrophages (nor neutrophils) during this particular type of infection. Alternatively, we attempted to administer liposome-encapsulated clodronate (Lipo-clodronate) as described previously (Nguyen-Chi et al., 2017; Phan et al., 2018; Travnickova et al., 2015). In our hands however, administration of 5 mg/ml Lipo-clodronate (3 nl) to 5 dpf larvae (instead of 2-3 dpf larvae), led to the rapid development of oedema.

Besides differences between the overall macrophage and neutrophil (inflammatory) response, the differential distribution of these cells was also investigated *in vivo* during infection utilising the transparency of the zebrafish and the availability of transgenic lines marking the vasculature. Neutrophils were never observed inside the cardinal caudal vein although in infected individuals they were certainly recruited and were observed in close contact with the outer vessel’s endothelium. Conversely, macrophages could be seen both outside and inside the vessel and the total proportion differed between high- and low-infected individuals. While in low-infected individuals the majority of macrophages recruited to the cardinal caudal vein remained outside the vessel in close contact with the endothelium, in high-infected individuals the majority of macrophages were recruited inside the caudal vein and were tightly attached to the luminal vessel wall. To our knowledge, such detailed description of the relative (re)distribution of neutrophils and macrophages, *in vivo*, during a trypanosome infection, has not been reported before.

Interestingly, exclusively in high-infected individuals, by 4 dpi large, round, dark and granular cells were observed, already under the bright field view, in the lumen of the cardinal caudal vein. These cells were confirmed to be foamy macrophages with high cytoplasmic lipid content. Foam cells, or foamy macrophages have been named after the lipid bodies accumulated in their cytoplasm leading to their typical enlarged morphological appearance (Dvorak et al., 1983), but are also distinguished by the presence of diverse cytoplasmic organelles (Melo et al., 2003). They have been associated with several (intracellular) infectious diseases, including Leishmaniasis, Chagas disease, experimental malaria, toxoplasmosis and tuberculosis, (reviewed in (López-Muñoz et al., 2018; Vallochi et al., 2018)) but never before with (extracellular) trypanosome infection. For example, during *T. cruzi* infection of rat, increased numbers of activated monocytes or macrophages were reported in the blood or heart (Melo and Machado, 2001). Interestingly, trypanosome uptake was shown to directly initiate the formation of lipid bodies in macrophages, leading to the appearance of foamy macrophages (D’Avila et al., 2011). During human *Mycobacterium tuberculosis* infections, foamy macrophages play a role in sustaining the presence of bacteria and contribute to tissue cavitation enabling the spread of the infection (Russell et al., 2009). Independently of the disease, it is clear that foamy macrophages are generally associated with inflammation, since their cytoplasmic lipid bodies are a source of eicosanoids, strong mediators of inflammation (Melo et al., 2006; Wymann and Schneiter, 2008). In turn, inflammatory mediators such as Prostaglandin E2 benefit trypanosome survival, as shown in *Trypanosoma*, *Leishmania*, *Plasmodium*, and *Toxoplasma* infections (reviewed in Vallochi et al., 2018). Our results are consistent with these reports as we show the occurrence of foamy macrophages exclusively in individuals that developed high parasitaemia, characterized by a strong pro-inflammatory response, and ultimately succumbed to the infection. Although we did not systematically investigate the exact kinetics of parasitaemia development in correlation with foamy macrophages occurrence, during our *in vivo* monitoring, we consistently observed that the increase in trypanosome number preceded the appearance of foamy macrophages. It is possible that, in high-infected individuals, foamy macrophages are formed due to the necessity to clear the increasing concentration of circulating trypanosome-derived moieties or of dying trypanosomes. The interaction with trypanosome-derived molecules, including soluble surface (glyco)proteins or trypanosome DNA, may not only be responsible for the activation of pro-inflammatory pathways, but also for a change in cell metabolism. The occurrence of foamy macrophages has been reported for intracellular trypanosomatids (*T. cruzi, Leishmania)*, and arachidonic acid-derived lipids were reported to act as regulators of the host immune response and trypanosome burden during *T. brucei* infections (López-Muñoz et al., 2018). To our knowledge our study is the first to report the presence of foamy macrophages during an extracellular trypanosome infection.

The possibility to detect the occurrence of large, granular cells already in the bright field and the availability of transgenic lines that allowed us to identify these cells as macrophages, further emphasizes the power of the zebrafish model. It allowed us to visualise *in vivo*, in real time, not only their occurrence but also their differential distribution with respect to other macrophages or neutrophils. Observations that we might have missed if we for example were to bleed an animal, perform immunohistochemistry or gene expression analysis. Thus, the possibility to separate high- and low-infected animals without the need to sacrifice them, allowed us to follow at the individual level the progression of the infection and the ensuing differential immune response.

In the future it will be interesting to analyse the transcription profiles of sorted macrophage populations from low- and high-infected larvae. Given the marked heterogeneity in macrophage activation observed especially within high-infected individuals, single-cell transcriptome analysis, of foamy macrophages in particular, may provide insights in the differential activation state of the various macrophage phenotypes. Furthermore, the zebrafish has already emerged as a valuable animal model to study inflammation and host-pathogen interaction and can be a powerful complementary tool to examine macrophage plasticity and polarization *in vivo*, by truly reflecting the complex nature of the environment during an ongoing infection in a live host. Finally, the availability of (partly) transparent adult zebrafish lines (Antinucci and Hindges, 2016; White et al., 2008), may aid the *in vivo* analysis of macrophage activation in adult individuals.

Altogether, in this study we describe the innate immune response of zebrafish larvae to *T. carassii* infection. The transparency and availability of various transgenic zebrafish lines, enabled us to establish a clinical scoring system that allowed us to monitor parasitaemia development and describe the differential response of neutrophils and macrophages at the individual level. Interestingly, for the first time in an extracellular trypanosome infection, we report the occurrence of foamy macrophages, characterized by a high lipid content and strong inflammatory profile, associated with susceptibility to the infection. Our model paves the way to investigate which mediators of the trypanosomes are responsible for the induction of such inflammatory response as well as study the condition that lead to the formation of foamy macrophages *in vivo*.

## Acknowledgments

This work was supported by the FishForPharma FP7 People: Marie Curie Initial Training Network, project number PITN-GA-2011-289209 and by the Dutch Research Council (NWO), project number 022.004.005. Dr. Christelle Langevin from Institut National de la Recherche Agronomique (INRA) is greatly acknowledged for her assistance with immunohistochemical analysis. The authors like to thank Dr. Danilo Pietretti, Marleen Scheer, and Dr. Sylvia Brugman from the Cell Biology and Immunology Group of Wageningen University & Research (WUR) for technical support with the RQ-PCR analysis, parasite isolation and for fruitful discussions; the CARUS Aquatic Research Facility of WUR is acknowledged for fish rearing and husbandry. Prof. Mark Carrington, Cambridge University, is acknowledged for the fruitful discussions and for revising the manuscript. Furthermore, the authors like to thank Dr. Norbert de Ruijter from the Wageningen Light Microscopy Centre, and the Montpellier Resources Imagerie facility for their assistance.

## Supplementary files

**S1 Fig.**
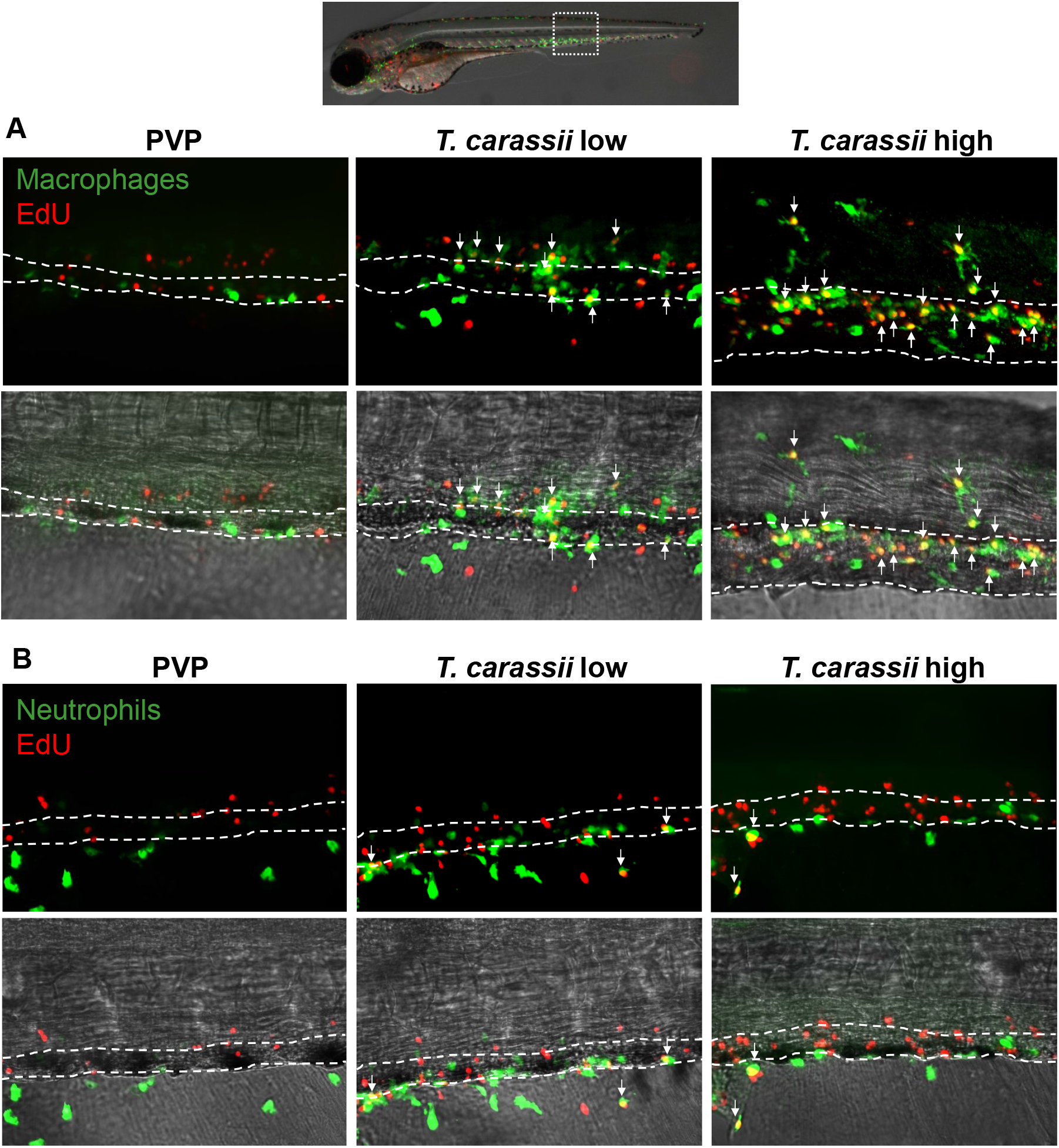
Differential distribution of EdU^+^ macrophages and neutrophils along the caudal vein of high- and low-infected larvae. Zebrafish were treated as described in figure 5. **A)** A high number of macrophages can be seen around and inside the caudal vein. Especially in high-infected individuals, the majority of cells within the vessel was EdU^+^, suggesting that in these larvae, proliferating macrophages migrated to the vessels. **B)** Neutrophils were never observed within the caudal vein and, independently of whether they proliferated (EdU^+^) or not, were mostly observed lining the vessel.

**S1 video.**
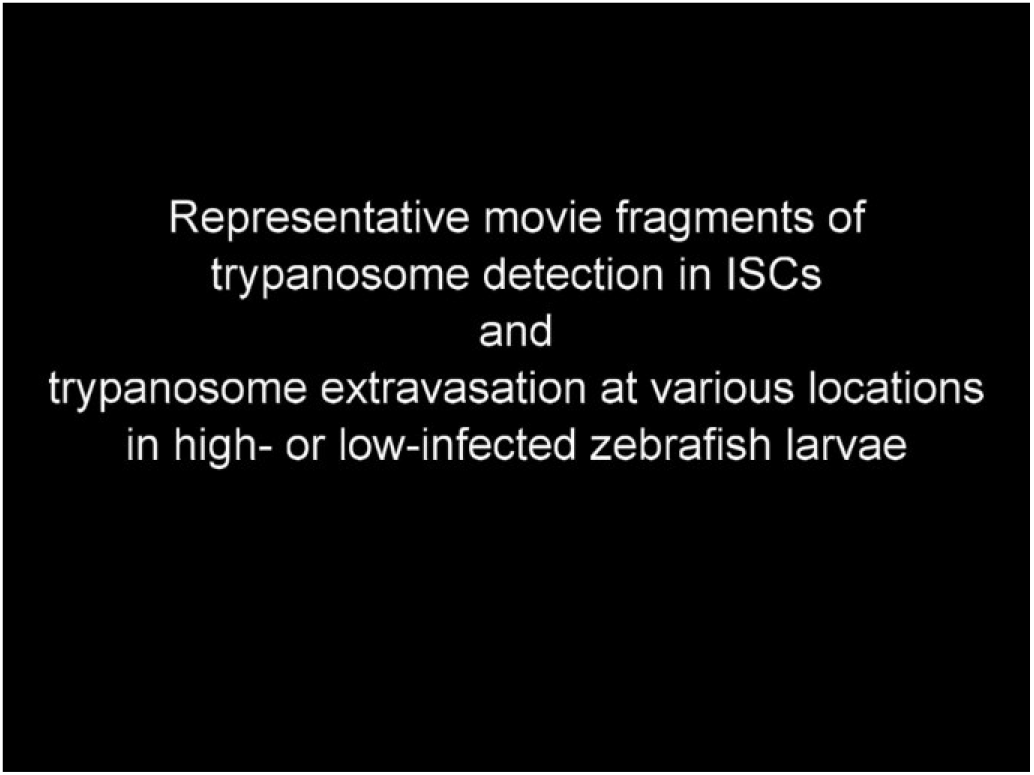
Clinical signs of *T. carassii* infection in high- and low-infected zebrafish larvae. *Tg(mpeg1.4:mCherry-F;mpx:GFP)* 5 dpf zebrafish were injected with *n*=200 *T. carassii* or with PVP and imaged at various time points after infection. Shown are high-speed videos (500 frames/sec, fps) or real-time videos (20 fps) capturing trypanosomes *in vivo* in blood or in tissues, as well as describing typical signs of anaemia and vasodilation.

**S2 video.**
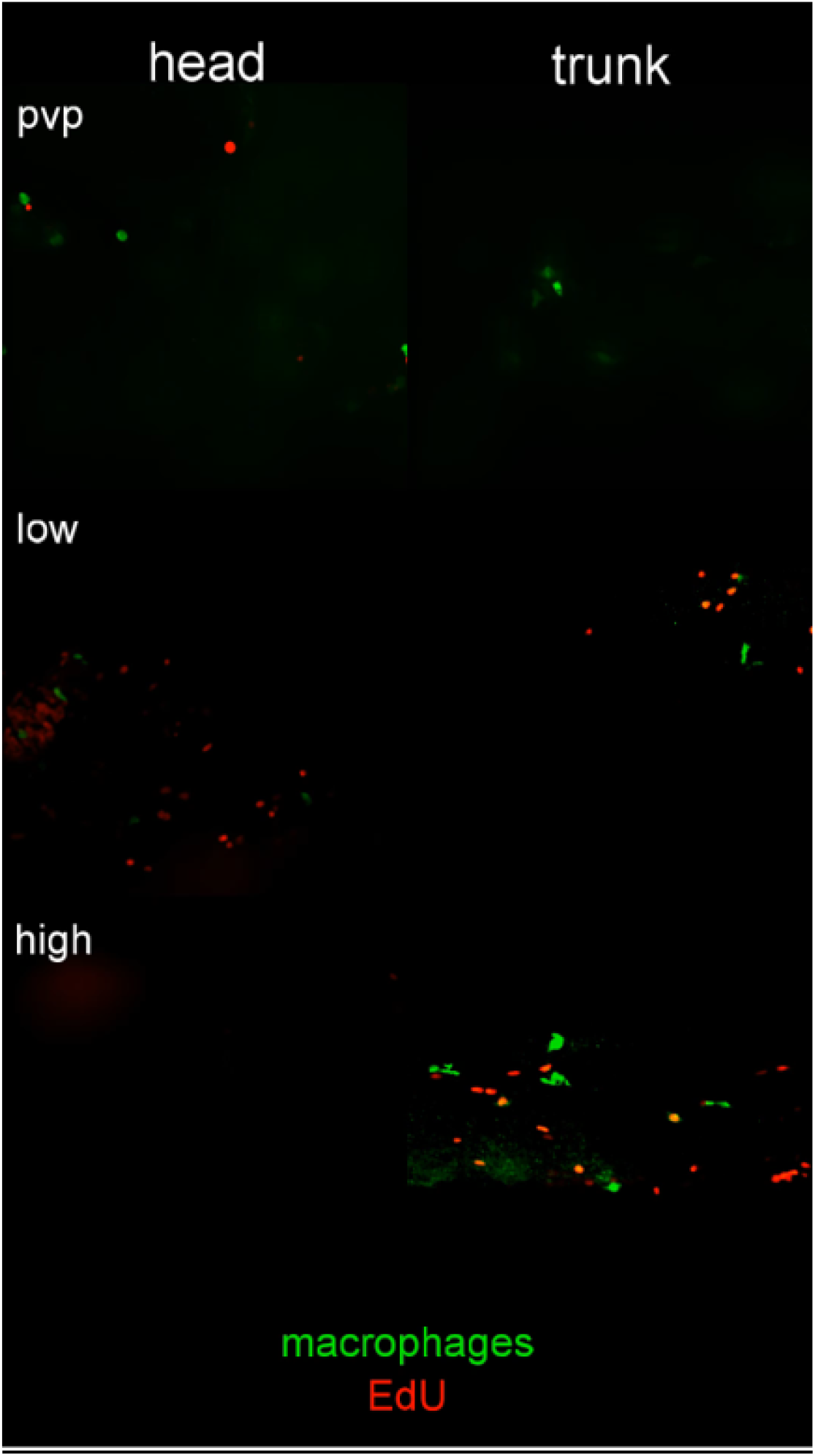
*T. carassii* infection triggers macrophage proliferation. AVI files corresponding to the maximum projection images shown in figure 5; Arrows are positioned as in figure 5, and indicate the location of EdU^+^ macrophages

**S3 video.**
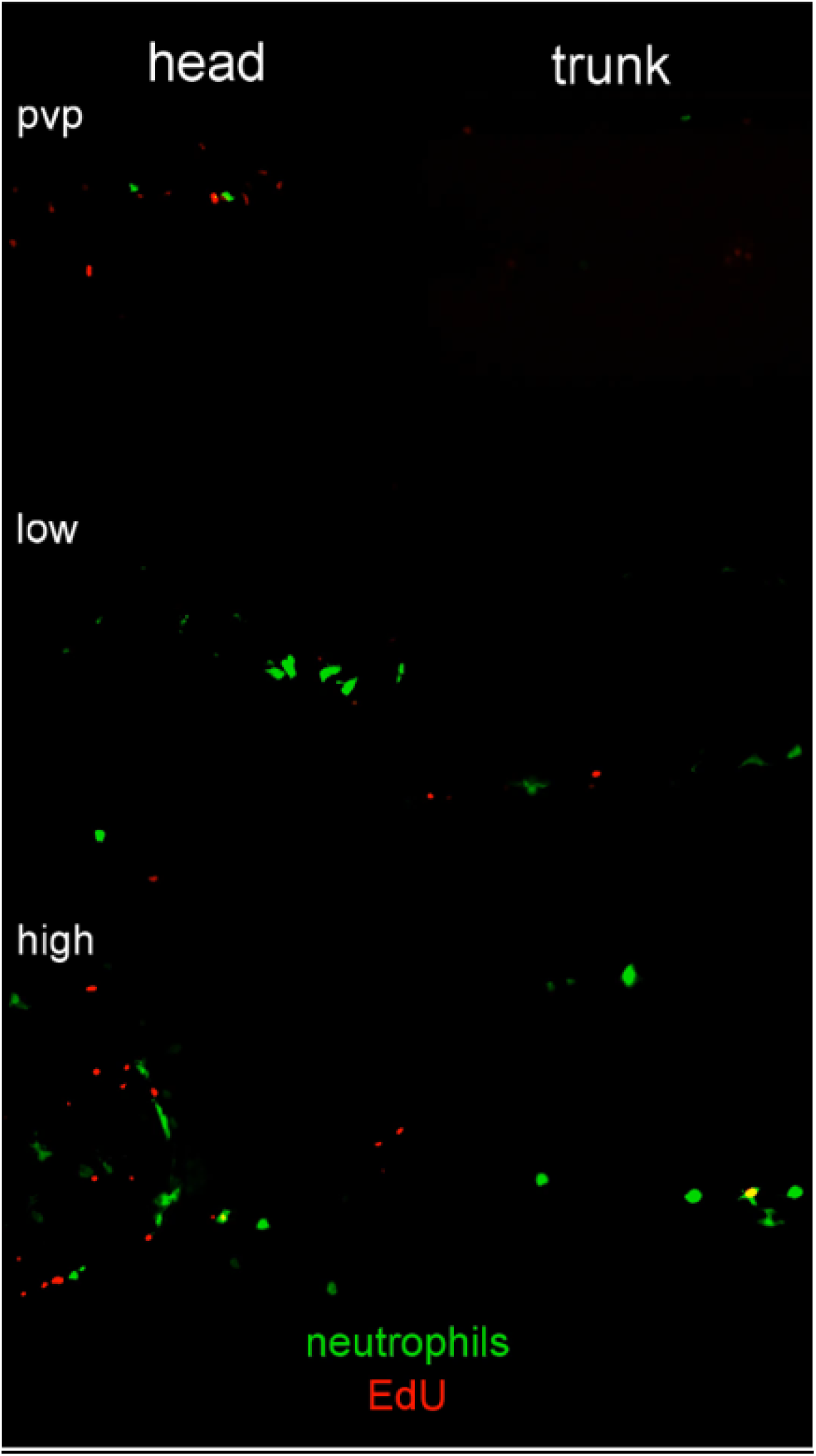
*T. carassii* infection triggers neutrophils proliferation. AVI files corresponding to the maximum projection images shown in figure 6; Arrows are positioned as in figure 6, and indicate the location of EdU^+^ neutrophils

**S4 video.**
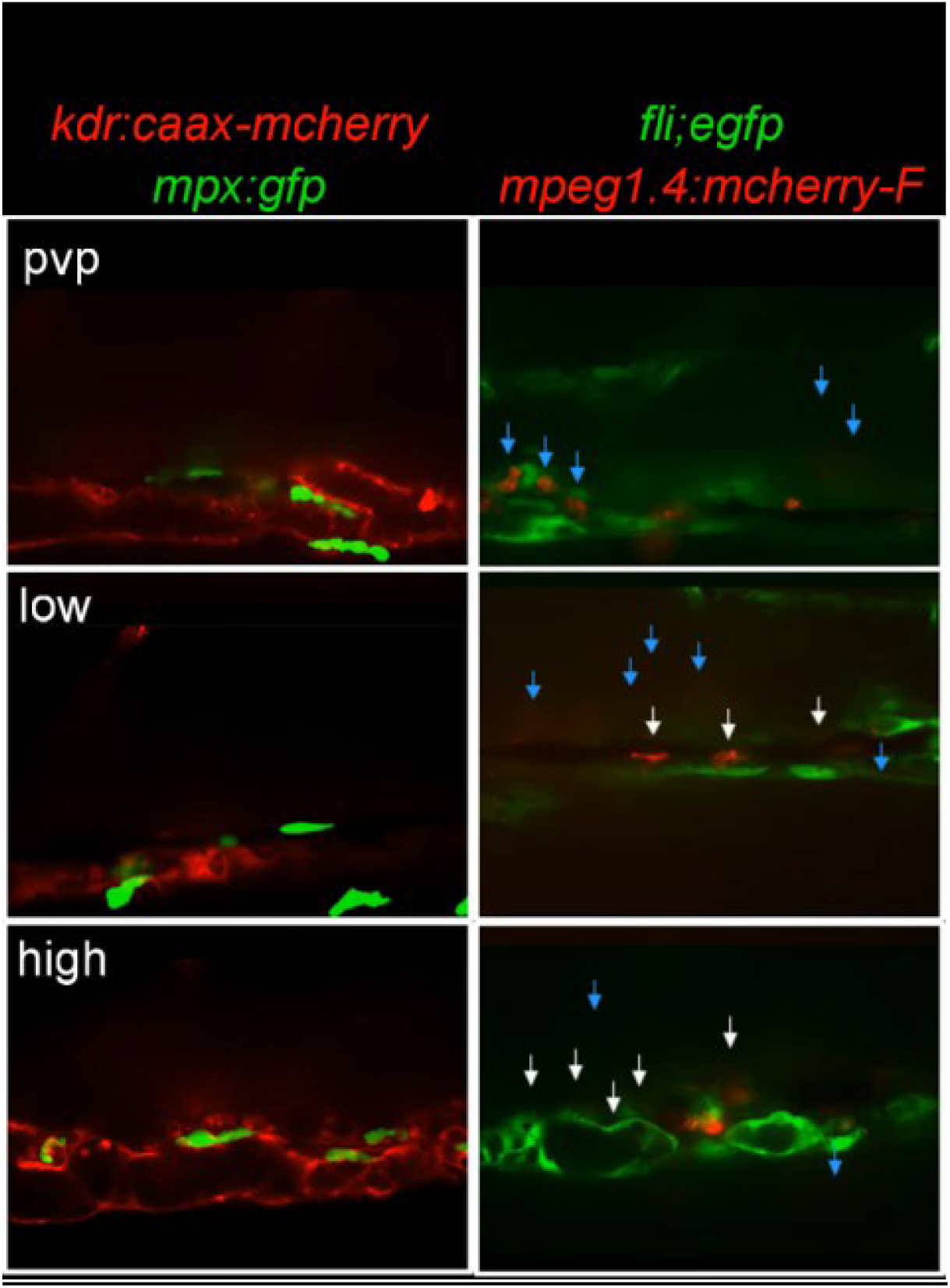
Macrophages are recruited into the cardinal caudal vein of high-infected zebrafish larvae. Zebrafish larvae were treated and imaged as described in figure 7. Shown are the AVI files corresponding to the maximum projection images shown in figure 7; Neutrophils were never observed within the vessel independently of the infection level (left panels). Macrophages however, could be seen outside (blue arrows) and inside the vessel (white arrows). The number of rounded macrophages inside the vessel increased with the infection level.

**S5 Video.**
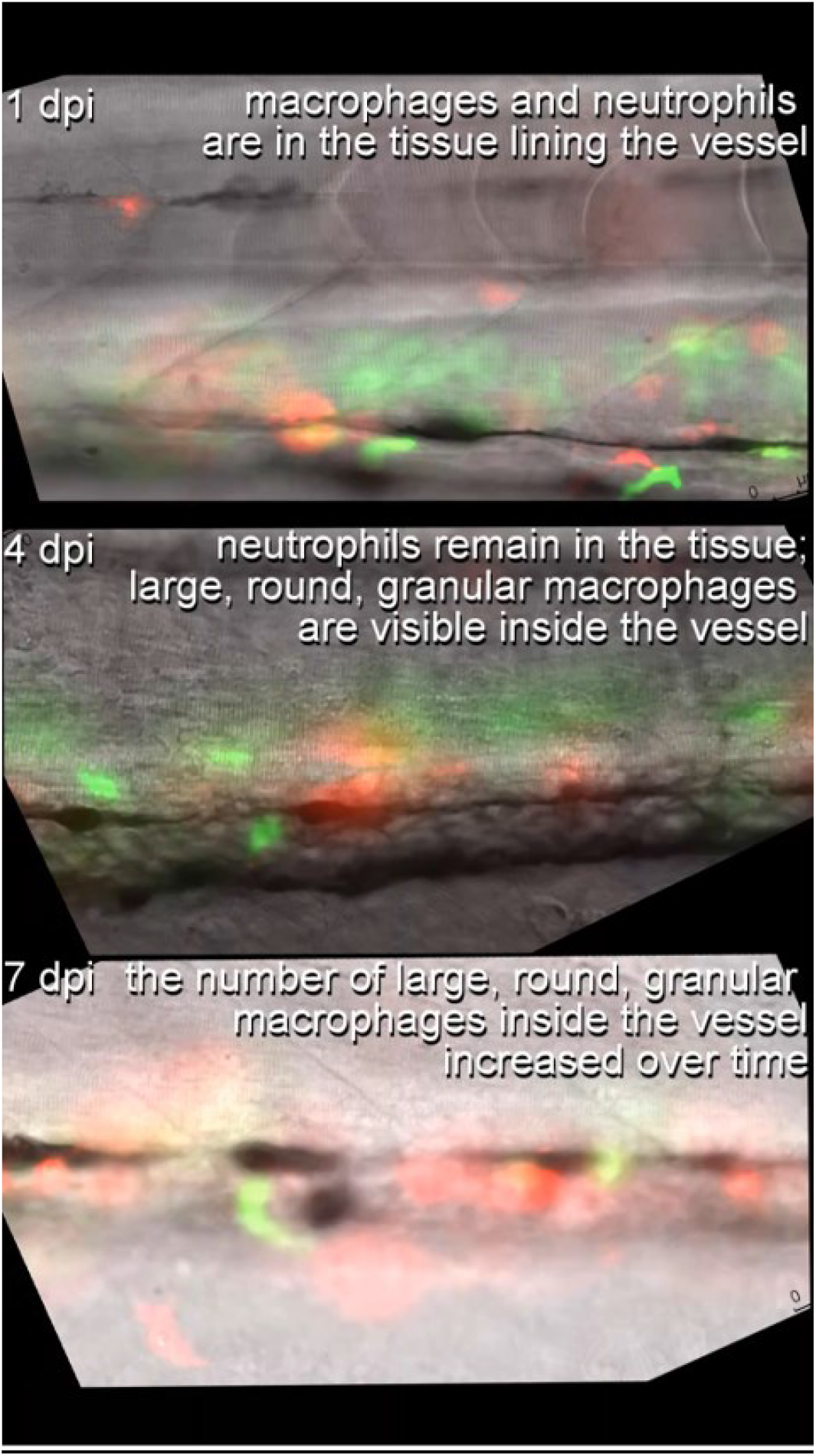
The occurrence of large granular macrophages increases with the progression of the infection in high-infected individuals. *Tg(mpeg1.4:mCherry-F;mpx:GFP)* zebrafish larvae were injected intravenously at 5 dpf with n=200 *T. carassii* or with PVP. At 4 dpi larvae were separated into high- and low-infected individuals and imaged at 5 dpi with a DMi8 inverted digital Leica microscope. The occurrence of large macrophages (arrows) in the cardinal caudal vessel increased with the progression of the infection and was exclusive to high infected individuals (4 and 7 dpi).

**S6 Video.**
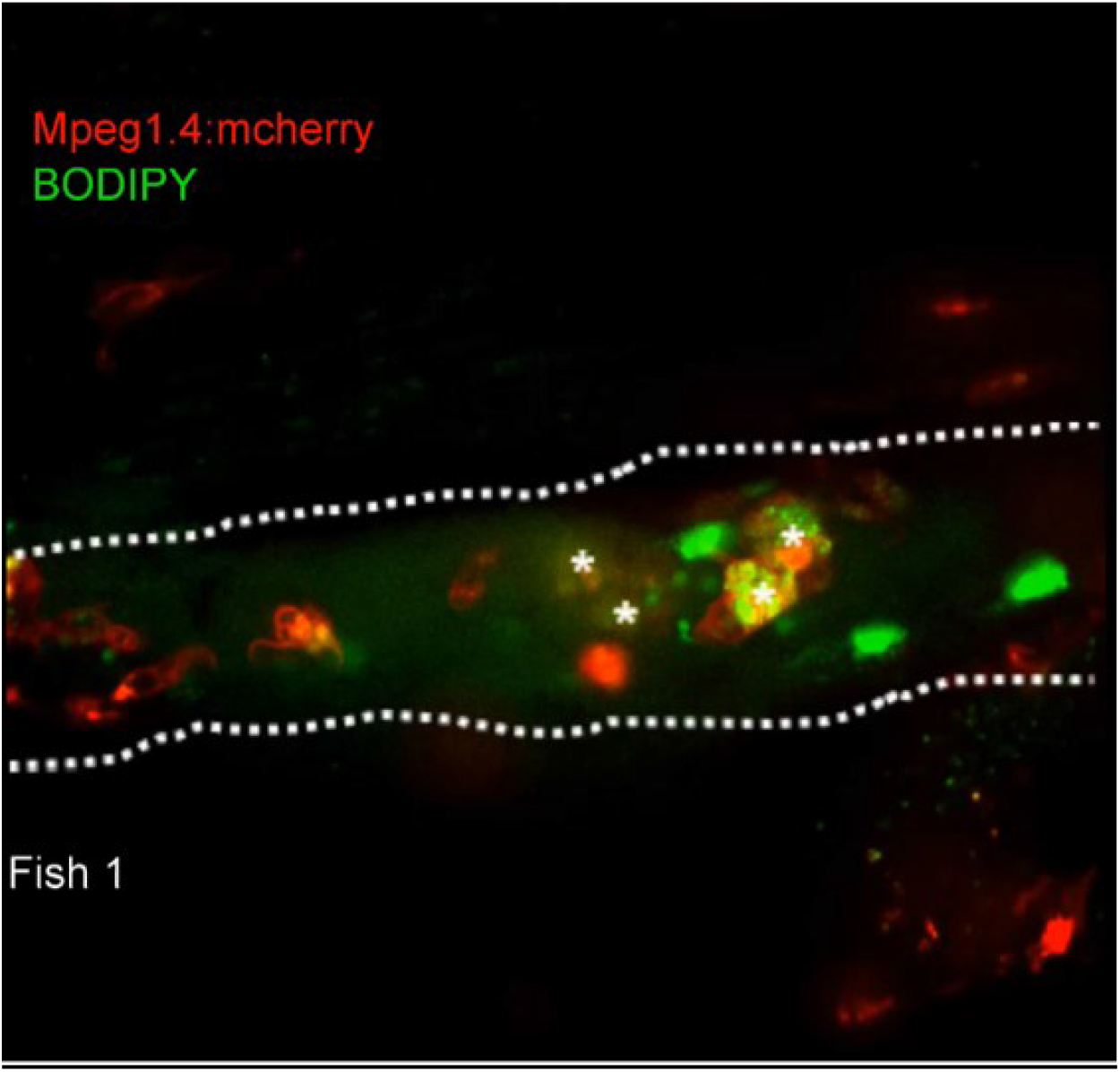
Large granular macrophages inside the vessel of high-infected larvae are rich in lipid bodies. *Tg(mpeg1.4:mCherry-F)* zebrafish larvae were infected intravenously at 5 dpf with n=200 *T. carassii* or with PVP. At 3 dpi, larvae received 1 nl of 30 μM BODIPY-FLC5 and were imaged 18-20 hours later using a Roper Spinning Disc Confocal Microscope at a 40x magnification. The AVI files corresponding to the maximum projection images shown in figure 8, as well as a second individual, are shown. Asterisks indicate the position of foamy macrophages inside the caudal vessel (dashed line).

## References

Antinucci P, Hindges R. 2016. A crystal-clear zebrafish for in vivo imaging. Sci Rep 6. doi:10.1038/srep29490

Baral TN, De Baetselier P, Brombacher F, Magez S. 2007. Control of Trypanosoma evansi Infection Is IgM Mediated and Does Not Require a Type I Inflammatory Response. J Infect Dis 195:1513–1520. doi:doi:10.1086/515577

Benard EL, Racz PI, Rougeot J, Nezhinsky AE, Verbeek FJ, Spaink HP, Meijer AH. 2015. Macrophage-expressed perforins Mpeg1 and Mpeg1.2 have an anti-bacterial function in zebrafish. J Innate Immun 7:136–152. doi:10.1159/000366103

Bertrand JY, Chi NC, Santoso B, Teng S, Stainier DYR, Traver D. 2010. Haematopoietic stem cells derive directly from aortic endothelium during development. Nature 464:108–111. doi:10.1038/nature08738

Boada-Sucre AA, Rossi Spadafora MS, Tavares-Marques LM, Finol HJ, Reyna-Bello A. 2016. Trypanosoma vivax Adhesion to Red Blood Cells in Experimentally Infected Sheep. Patholog Res Int 2016. doi:10.1155/2016/4503214

Caljon G, Mabille D, Stijlemans B, Trez C De, Mazzone M, Tacchini-cottier F, Malissen M, Ginderachter JA Van, Magez S, Baetselier P De, Abbeele J Van Den. 2018. Neutrophils enhance early Trypanosoma brucei infection onset 1–11. doi:10.1038/s41598-018-29527-y

Chi NC, Shaw RM, Val S De, Kang G, Jan LY, Black BL, Stainier DYR. 2008. Expression and Atrioventricular Canal Formation. Genes Dev 734–739. doi:10.1101/gad.1629408.734

Cnops J, De Trez C, Stijlemans B, Keirsse J, Kauffmann F, Barkhuizen M, Keeton R, Boon L, Brombacher F, Magez S. 2015. NK-, NKT- and CD8-Derived IFNγ Drives Myeloid Cell Activation and Erythrophagocytosis, Resulting in Trypanosomosis-Associated Acute Anemia. PLoS Pathog 11. doi:10.1371/journal.ppat.1004964

Coller SP, Mansfield JM, Paulnock DM. 2003. Glycosylinositolphosphate Soluble Variant Surface Glycoprotein Inhibits IFN-γ-Induced Nitric Oxide Production Via Reduction in STAT1 Phosphorylation in African Trypanosomiasis. J Immunol 171:1466–1472. doi:10.4049/jimmunol.171.3.1466

Cronan MR, Tobin DM. 2014. Fit for consumption: zebrafish as a model for tuberculosis. Dis Model Mech. doi:10.1242/dmm.016089

D’Avila H, Freire-de-Lima CG, Roque NR, Teixeira L, Barja-Fidalgo C, Silva AR, Melo RCN, DosReis GA, Castro-Faria-Neto HC, Bozza PT. 2011. Host cell lipid bodies triggered by Trypanosoma cruzi infection and enhanced by the uptake of apoptotic cells are associated with prostaglandin E2 generation and increased parasite growth. J Infect Dis 204:951–961. doi:10.1093/infdis/jir432

Daulouede S, Bouteille B, Moynet D, De Baetselier P, Courtois P, Lemesre JL, Buguet A, Cespuglio R, Vincendeau P. 2001. Human macrophage tumor necrosis factor (TNF)-alpha production induced by Trypanosoma brucei gambiense and the role of TNF-alpha in parasite control. J Infect Dis 183:988–991.

Davison JM, Akitake CM, Goll MG, Rhee JM, Gosse N, Baier H, Halpern ME, Leach SD, Parsons MJ. 2007. Transactivation from Gal4-VP16 transgenic insertions for tissue-specific cell labeling and ablation in zebrafish. Dev Biol 304:811–824. doi:10.1016/j.ydbio.2007.01.033

Dóró É, Jacobs SH, Hammond FR, Schipper H, Pieters RP, Carrington M, Wiegertjes GF, Forlenza M. 2019. Visualizing trypanosomes in a vertebrate host reveals novel swimming behaviours, adaptations and attachment mechanisms. Elife 8. doi:10.7554/elife.48388

Dvorak AM, Dvorak HF, Peters SP, Shulman ES, MacGlashan DWJ, Pyne K, Harvey VS, Galli SJ, Lichtenstein LM. 1983. Lipid bodies: cytoplasmic organelles important to arachidonate metabolism in macrophages and mast cells. J Immunol 131:2965–2976.

Ellett F, Pase L, Hayman JW, Andrianopoulos A, Lieschke GJ. 2011. Mpeg1 Promoter Transgenes Direct Macrophage-Lineage Expression in Zebrafish. Blood 117:e49–e56. doi:10.1182/blood-2010-10-314120

Engstler M, Pfohl T, Herminghaus S, Boshart M, Wiegertjes G, Heddergott N, Overath P. 2007. Hydrodynamic Flow-Mediated Protein Sorting on the Cell Surface of Trypanosomes. Cell 131:505–515. doi:10.1016/j.cell.2007.08.046

Forlenza M, Kaiser T, Savelkoul HFJ, Wiegertjes GF. 2012. The use of real-time quantitative PCR for the analysis of cytokine mRNA levels.Methods in Molecular Biology (Clifton, N.J.). pp. 7–23. doi:10.1007/978-1-61779-439-1_2

Forlenza M, Magez S, Scharsack JP, Westphal A, Savelkoul HFJ, Wiegertjes GF. 2009. Receptor-Mediated and Lectin-Like Activities of Carp (Cyprinus carpio) TNF-α. J Immunol 183:5319–5332. doi:10.4049/jimmunol.0901780

García-Valtanen P, Martínez-López A, López-Muñoz A, Bello-Perez M, Medina-Gali RM, Ortega-Villaizán MDM, Varela M, Figueras A, Mulero V, Novoa B, Estepa A, Coll J. 2017. Zebra fish lacking adaptive immunity acquire an antiviral alert state characterized by upregulated gene expression of apoptosis, multigene families, and interferon-related genes. Front Immunol 8:121. doi:10.3389/fimmu.2017.00121

Guegan F, Plazolles N, Baltz T, Coustou V. 2013. Erythrophagocytosis of desialylated red blood cells is responsible for anaemia during Trypanosomavivax infection. Cell Microbiol 15:1285–1303. doi:10.1111/cmi.12123

Iraqi F, Sekikawa K, Rowlands J, Teale A. 2001. Susceptibility of tumour necrosis factor-alpha genetically deficient mice to Trypanosoma congolense infection. Parasite Immunol 23:445–451.

Islam A, Woo P. 1991. Anemia and its mechanism in goldfish Carassius auratus infected with Trypanosoma danilewskyi. Dis Aquat Organ 11:37–43. doi:10.3354/dao011037

Joerink M, Groeneveld A, Ducro B, Savelkoul HFJ, Wiegertjes GF. 2007. Mixed infection with Trypanoplasma borreli and Trypanosoma carassii induces protection: Involvement of cross-reactive antibodies. Dev Comp Immunol 31:903–915. doi:10.1016/j.dci.2006.12.003

Joerink M, Saeij J, Stafford J, Belosevic M, Wiegertjes G. 2004. Animal models for the study of innate immunity: protozoan infections in fish In: Flik G, Wiegertjes GF, editors. Host-Parasite Interactions. Taylor & Francis. p. (55):67–89.

Kaushik RS, Uzonna JE, Gordon JR, Tabel H. 1999. Innate resistance to Trypanosoma congolense infections: Differential production of nitric oxide by macrophages from susceptible BALB/c and resistant C57B1/6 mice. Exp Parasitol 92:131–143. doi:10.1006/expr.1999.4408

Kent M, Lom J, Dykova I, Dyková I. 1993. Protozoan Parasites of Fishes. J Parasitol 79:673. doi:10.2307/3283600

Kovacevic N, Hagen MO, Xie J, Belosevic M. 2015. The analysis of the acute phase response during the course of Trypanosoma carassii infection in the goldfish (Carassius auratus L.). Dev Comp Immunol 53:112–122. doi:10.1016/j.dci.2015.06.009

Krettli AU, Weisz-Carrington P, Nussenzweig RS. 1979. Membrane-bound antibodies to bloodstream Trypanosoma cruzi in mice: Strain differences in susceptibility to complement-mediated lysis. Clin Exp Immunol 37:416–423.

La Greca F, Haynes C, Stijlemans B, De Trez C, Magez S. 2014. Antibody-mediated control of Trypanosoma vivax infection fails in the absence of tumour necrosis factor. Parasite Immunol 36:271–276. doi:10.1111/pim.12106

Langenau DM, Ferrando AA, Traver D, Kutok JL, Hezel JP, Kanki JP, Zon LI, Look AT, Trede NS. 2004. In vivo tracking of T cell development, ablation, and engraftment in transgenic zebrafish. Proc Natl Acad Sci U S A 101:7369–7374.

Lawson ND, Weinstein BM. 2002. In vivo imaging of embryonic vascular development using transgenic zebrafish. Dev Biol 248:307–318. doi:10.1006/dbio.2002.0711

Lischke A, Klein C, Stierhof Y-D, Hempel M, Mehlert A, Almeida IC, Ferguson MAJ, Overath P. 2000. Isolation and characterization of glycosylphosphatidylinositol-anchored, mucin-like surface glycoproteins from bloodstream forms of the freshwater-fish parasite Trypanosoma carassii. Biochem J 345:693. doi:10.1042/0264-6021:3450693

López-Muñoz RA, Molina-Berríos A, Campos-Estrada C, Abarca-Sanhueza P, Urrutia-Llancaqueo L, Peña-Espinoza M, Maya JD. 2018. Inflammatory and Pro-resolving Lipids in Trypanosomatid Infections: A Key to Understanding Parasite Control. Front Microbiol 9:1961. doi:10.3389/fmicb.2018.01961

Lopez R, Demick KP, Mansfield JM, Paulnock DM. 2008. Type I IFNs Play a Role in Early Resistance, but Subsequent Susceptibility, to the African Trypanosomes. J Immunol 181:4908–4917. doi:10.4049/jimmunol.181.7.4908

Lucas Rudolf, Magez S, De Leys R, Fransen L, Scheerlinck JP, Rampelberg M, Sablon E, De Baetselier P. 1994. Mapping the lectin-like activity of tumor necrosis factor. Science (80-) 263:814–817. doi:10.1126/science.8303299

Lucas R, Magez S, De Leys R, Fransen L, Scheerlinck JP, Rampelberg M, Sablon E, De Baetselier P. 1994. Mapping the lectin-like activity of tumor necrosis factor. Science (80-) 263:814–817.

Magez S., Lucas R, Darji A, Bajyana Songa E, Hamers R, Baetselier P de. 1993. Murine tumour necrosis factor plays a protective role during the initial phase of the experimental infection with Trypanosoma brucei brucei. Parasite Immunol 15:635–641. doi:10.1111/j.1365-3024.1993.tb00577.x

Magez S, Caljon G. 2011. Mouse models for pathogenic African trypanosomes: unravelling the immunology of host-parasite-vector interactions. Parasite Immunol 33:423–429. doi:10.1111/j.1365-3024.2011.01293.x

Magez S, Geuskens M, Beschin A, del Favero H, Verschueren H, Lucas R, Pays E, de Baetselier P. 1997. Specific uptake of tumor necrosis factor-alpha is involved in growth control of Trypanosoma brucei. J Cell Biol 137:715–727.

Magez S, Radwanska M, Beschin A, Sekikawa K, De Baetselier P. 1999. Tumor necrosis factor alpha is a key mediator in the regulation of experimental Trypanosoma brucei infections. Infect Immun 67:3128–3132.

Magez S, Radwanska M, Drennan M, Fick L, Baral TN, Allie N, Jacobs M, Nedospasov S, Brombacher F, Ryffel B, Baetselier P De. 2007. Tumor Necrosis Factor (TNF) Receptor–1 (TNFp55) Signal Transduction and Macrophage-Derived Soluble TNF Are Crucial for Nitric Oxide–Mediated Trypanosoma congolense Parasite Killing. J Infect Dis 196:954–962. doi:10.1086/520815

Magez S, Radwanska M, Drennan M, Fick L, Baral TN, Brombacher F, De Baetselier P. 2006. Interferon-gamma and nitric oxide in combination with antibodies are key protective host immune factors during Trypanosoma congolense Tc13 Infections. J Infect Dis 193:1575–1583.

Magez S, Radwanska M, Stijlemans B, Xong H V, Pays E, De Baetselier P. 2001. A conserved flagellar pocket exposed high mannose moiety is used by African trypanosomes as a host cytokine binding molecule. J Biol Chem 276:33458–33464.

Magez S, Stijlemans B, Baral T, De Baetselier P. 2002. VSG-GPI anchors of African trypanosomes: their role in macrophage activation and induction of infection-associated immunopathology. Microbes Infect 4:999–1006. doi:10.1016/S1286-4579(02)01617-9

Magez S, Stijlemans B, De Baetselier P, Radwanska M, Pays E, Ferguson MAJ. 1998. The glycosyl-inositol-phosphate and dimyristoylglycerol moieties of the glycosylphosphatidylinositol anchor of the trypanosome variant-specific surface glycoprotein are distinct macrophage-activating factors. J Immunol.

Mansfield JM, Paulnock DM. 2005. Regulation of innate and acquired immunity in African trypanosomiasis. Parasite Immunol 27:361–371. doi:10.1111/j.1365-3024.2005.00791.x

McAllister M, Phillips N, Belosevic M. 2019. Trypanosoma carassii infection in goldfish (Carassius auratus L.): changes in the expression of erythropoiesis and anemia regulatory genes. Parasitol Res 118:1147–1158. doi:10.1007/s00436-019-06246-5

Melo RC., Machado CR. 2001. Trypanosoma cruzi: Peripheral Blood Monocytes and Heart Macrophages in the Resistance to Acute Experimental Infection in Rats. Exp Parasitol 97:15–23. doi:10.1006/EXPR.2000.4576

Melo RCN, D’Ávila H, Fabrino DL, Almeida PE, Bozza PT. 2003. Macrophage lipid body induction by Chagas disease in vivo: putative intracellular domains for eicosanoid formation during infection. Tissue Cell 35:59–67. doi:10.1016/S0040-8166(02)00105-2

Melo RCN, Fabrino DL, Dias FF, Parreira GG. 2006. Lipid bodies: Structural markers of inflammatory macrophages in innate immunity. Inflamm Res 55:342–348. doi:10.1007/s00011-006-5205-0

Mishra RR, Senapati SK, Sahoo SC. 2017. Trypanosomiasis induced oxidative stress and hemato-biochemical alteration in cattle. Artic J Entomol Zool Stud 5:721–727.

Musoke AJ, Barbet AF. 1977. Activation of complement by variant-specific surface antigen of Trypanosoma brucei. Nature 270:438–440.

Naessens J. 2006. Bovine trypanotolerance: A natural ability to prevent severe anaemia and haemophagocytic syndrome? Int J Parasitol 36:521–528. doi:10.1016/j.ijpara.2006.02.012

Namangala B, De Baetselier P, Beschin A. 2009. Both Type-I and Type-II Responses Contribute to Murine Trypanotolerance. J Vet Med Sci 71:313–318.

Namangala B, Noel W, De Baetselier P, Brys L, Beschin A. 2001. Relative contribution of interferon-gamma and interleukin-10 to resistance to murine African trypanosomosis. J Infect Dis 183:1794–1800.

Nguyen-Chi M, Laplace-Builhé B, Travnickova J, Luz-Crawford P, Tejedor G, Lutfalla G, Kissa K, Jorgensen C, Djouad F. 2017. TNF signaling and macrophages govern fin regeneration in zebrafish larvae. Cell Death Dis 8:e2979–e2979. doi:10.1038/cddis.2017.374

Nguyen-Chi M, Laplace-Builhe B, Travnickova J, Luz-Crawford P, Tejedor G, Phan QT, Duroux-Richard I, Levraud J-P, Kissa K, Lutfalla G, Jorgensen C, Djouad F. 2015. Identification of polarized macrophage subsets in zebrafish. Elife 4:e07288. doi:10.7554/eLife.07288

Nguyen-Chi M, Phan QT, Gonzalez C, Dubremetz J-F, Levraud J-P, Lutfalla G. 2014a. Transient infection of the zebrafish notochord with E. coli induces chronic inflammation. Dis Model Mech 7:871–82. doi:10.1242/dmm.014498

Nguyen-Chi M, Phan QT, Gonzalez C, Dubremetz J-F, Levraud J-P, Lutfalla G. 2014b. Transient infection of the zebrafish notochord with E. coli induces chronic inflammation. Dis Model Mech 7:871–82. doi:10.1242/dmm.014498

Noël W, Hassanzadeh G, Raes G, Namangala B, Daems I, Brys L, Brombacher F, Baetselier PD, Beschin A. 2002. Infection stage-dependent modulation of macrophage activation in Trypanosoma congolense-resistant and -susceptible mice. Infect Immun 70:6180–6187.

Noël W, Raes G, Ghassabeh GH, De Baetselier P, Beschin A. 2004. Alternatively activated macrophages during parasite infections. Trends Parasitol 20:126–133. doi:10.1016/j.pt.2004.01.004

O’Gorman GM, Park SDE, Hill EW, Meade KG, Mitchell LC, Agaba M, Gibson JP, Hanotte O, Naessens J, Kemp SJ, MacHugh DE. 2006. Cytokine mRNA profiling of peripheral blood mononuclear cells from trypanotolerant and trypanosusceptible cattle infected with Trypanosoma congolense. Physiol Genomics 28:53–61. doi:10.1152/physiolgenomics.00100.2006

Oladiran A, Beauparlant D, Belosevic M. 2011. The expression analysis of inflammatory and antimicrobial genes in the goldfish (Carassius auratus L.) infected with Trypanosoma carassii. Fish Shellfish Immunol 31:606–613. doi:10.1016/j.fsi.2011.07.008

Oladiran A, Belosevic M. 2012. Recombinant glycoprotein 63 (Gp63) of Trypanosoma carassii suppresses antimicrobial responses of goldfish (Carassius auratus L.) monocytes and macrophages q. Int J Parasitol 42:621–633. doi:10.1016/j.ijpara.2012.04.012

Oladiran A, Belosevic M. 2010. Trypanosoma carassii calreticulin binds host complement component C1q and inhibits classical complement pathway-mediated lysis. Dev Comp Immunol 34:396–405. doi:10.1016/j.dci.2009.11.005

Oladiran A, Belosevic M. 2009. Trypanosoma carassii hsp70 increases expression of inflammatory cytokines and chemokines in macrophages of the goldfish (Carassius auratus L.). Dev Comp Immunol 33:1128–1136. doi:10.1016/j.dci.2009.06.003

Overath P, Haag J, Lischke A, O’HUigin C. 2001. The surface structure of trypanosomes in relation to their molecular phylogeny. Int J Parasitol 31:468–471.

Overath P, Ruoff J, Stierhof YD, Haag J, Tichy H, Dyková I, Lom J. 1998. Cultivation of bloodstream forms of Trypanosoma carassii, a common parasite of freshwater fish. Parasitol Res 84:343–347. doi:10.1007/s004360050408

Page DM, Wittamer V, Bertrand JY, Lewis KL, Pratt DN, Delgado N, Schale SE, McGue C, Jacobsen BH, Doty A, Pao Y, Yang H, Chi NC, Magor BG, Traver D. 2013. An evolutionarily conserved program of B-cell development and activation in zebrafish. Blood 122:e1–e11. doi:10.1182/blood-2012-12-471029

Palha N, Guivel-Benhassine F, Briolat V, Lutfalla G, Sourisseau M, Ellett F, Wang CH, Lieschke GJ, Herbomel P, Schwartz O, Levraud JP. 2013. Real-Time Whole-Body Visualization of Chikungunya Virus Infection and Host Interferon Response in Zebrafish. PLoS Pathog 9:e1003619. doi:10.1371/journal.ppat.1003619

Petrie-Hanson L, Hohn C, Hanson L. 2009. Characterization of rag1 mutant zebrafish leukocytes. BMC Immunol 10:8. doi:10.1186/1471-2172-10-8

Phan QT, Sipka T, Gonzalez C, Levraud JP, Lutfalla G, Nguyen-Chi M. 2018. Neutrophils use superoxide to control bacterial infection at a distance. PLoS Pathog 14:1–29. doi:10.1371/journal.ppat.1007157

Radwanska M, Vereecke N, Deleeuw V, Pinto J, Magez S. 2018. Salivarian Trypanosomosis: A Review of Parasites Involved, Their Global Distribution and Their Interaction With the Innate and Adaptive Mammalian Host Immune System. Front Immunol 9:1–20. doi:10.3389/fimmu.2018.02253

Ramakrishnan L. 2013. The zebrafish guide to tuberculosis immunity and treatment. Cold Spring Harb Symp Quant Biol 78:179–192. doi:10.1101/sqb.2013.78.023283

Renshaw SA, Loynes CA, Trushell DMI, Elworthy S, Ingham PW, Whyte MKB. 2006. A transgenic zebrafish model of neutrophilic inflammation. Blood 108:3976–3978. doi:10.1182/blood-2006-05-024075

Renshaw SA, Trede NS. 2012. A model 450 million years in the making: zebrafish and vertebrate immunity. Dis Model Mech 5:38–47. doi:10.1242/dmm.007138

Ribeiro CMS, Pontes MJSL, Bird S, Chadzinska M, Scheer M, Verburg-van Kemenade BML, Savelkoul HFJ, Wiegertjes GF. 2010. Trypanosomiasis-induced Th17-like immune responses in carp. PLoS One 5:e13012. doi:10.1371/journal.pone.0013012

Rifkin MR, Landsberger FR. 1990. Trypanosome variant surface glycoprotein transfer to target membranes: A model for the pathogenesis of trypanosomiasis. Proc Natl Acad Sci U S A 87:801–805. doi:10.1073/pnas.87.2.801

Rosowski EE, Knox BP, Archambault LS, Huttenlocher A, Keller NP, Wheeler RT, Davis JM. 2018. The zebrafish as a model host for invasive fungal infections. J Fungi 4. doi:10.3390/jof4040136

Russell DG, Cardona P-J, Kim M-J, Allain S, Altare F. 2009. Foamy macrophages and the progression of the human tuberculosis granuloma. Nat Immunol 10:943–948. doi:10.1038/ni.1781

Saeij JPJ, Groeneveld A, Van Rooijen N, Haenen OLM, Wiegertjes GF. 2003. Minor effect of depletion of resident macrophages from peritoneal cavity on resistance of common carp Cyprinus carpio to blood flagellates. Dis Aquat Organ 57:67–75. doi:10.3354/dao057067

Scharsack JP, Steinhagen D, Kleczka C, Schmidt JO, Körting W, Michael RD, Leibold W, Schuberth HJ. 2003. Head kidney neutrophils of carp (Cyprinus carpio L.) are functionally modulated by the haemoflagellate Trypanoplasma borreli. Fish Shellfish Immunol 14:389–403. doi:10.1006/fsim.2002.0447

Sileghem M, Saya R, Grab DJ, Naessens J. 2001. An accessory role for the diacylglycerol moiety of variable surface glycoprotein of African trypanosomes in the stimulation of bovine monocytes. Vet Immunol Immunopathol 78:325–339. doi:10.1016/S0165-2427(01)00241-0

Simpson AGB, Stevens JR, Lukes J. 2006. The evolution and diversity of kinetoplastid flagellates. Trends Parasitol 22:168–174.

Sternberg JM, Mabbott NA. 1996. Nitric oxide-mediated suppression of T cell responses during Trypanosoma brucei infection: Soluble trypanosome products and interferon-γ are synergistic inducers of nitric oxide synthase. Eur J Immunol. doi:10.1002/eji.1830260306

Stijlemans B, Caljon G, Van Den Abbeele J, Van Ginderachter JA, Magez S, De Trez C. 2016. Immune evasion strategies of Trypanosoma brucei within the mammalian host: Progression to pathogenicity. Front Immunol. doi:10.3389/fimmu.2016.00233

Stijlemans B, Guilliams M, Raes G, Beschin A, Magez S, De Baetselier P. 2007. African trypanosomosis: from immune escape and immunopathology to immune intervention. Vet Parasitol 148:3–13.

Stijlemans B, Radwanska M, Trez C De, Magez S. 2017. African trypanosomes undermine humoral responses and vaccine development: Link with inflammatory responses? Front Immunol 8:582. doi:10.3389/fimmu.2017.00582

Torraca V, Masud S, Spaink HP, Meijer AH. 2014. Macrophage-pathogen interactions in infectious diseases: new therapeutic insights from the zebrafish host model. Dis Model Mech 7:785–797. doi:10.1242/dmm.015594

Torraca V, Mostowy S. 2018. Zebrafish Infection: From Pathogenesis to Cell Biology. Trends Cell Biol 28:143–156. doi:10.1016/j.tcb.2017.10.002

Travnickova J, Tran Chau V, Julien E, Mateos-Langerak J, Gonzalez C, Lelievre E, Lutfalla G, Tavian M, Kissa K. 2015. Primitive macrophages control HSPC mobilization and definitive haematopoiesis. Nat Commun 6:6227. doi:10.1038/ncomms7227

Vallochi AL, Teixeira L, Oliveira K da S, Maya-Monteiro CM, Bozza PT. 2018. Lipid Droplet, a Key Player in Host-Parasite Interactions. Front Immunol 9:1022. doi:10.3389/fimmu.2018.01022

White RM, Sessa A, Burke C, Bowman T, LeBlanc J, Ceol C, Bourque C, Dovey M, Goessling W, Burns CE, Zon LI. 2008. Transparent Adult Zebrafish as a Tool for In Vivo Transplantation Analysis. Cell Stem Cell. doi:10.1016/j.stem.2007.11.002

Woo PTK, Ardelli BF. 2014. Immunity against selected piscine flagellates. Dev Comp Immunol 43:268–279. doi:10.1016/j.dci.2013.07.006

Wu H, Liu G, Shi M. 2017. Interferon gamma in African trypanosome infections: Friends or foes? Front Immunol. doi:10.3389/fimmu.2017.01105

Wymann MP, Schneiter R. 2008. Lipid signalling in disease. Nat Rev Mol Cell Biol. doi:10.1038/nrm2335

